# Rapid compensatory evolution can rescue low fitness symbioses following partner-switching

**DOI:** 10.1101/2020.11.06.371401

**Authors:** Megan E S Sørensen, A Jamie Wood, Duncan D Cameron, Michael A Brockhurst

## Abstract

Partner-switching plays an important role in the evolution of symbiosis, enabling local adaptation and recovery from the breakdown of symbiosis. Because of intergenomic epistasis, partner-switched symbioses may possess novel combinations of phenotypes but may also exhibit low fitness due to their lack of recent coevolutionary history. Here, we examine the structure and mechanisms of intergenomic epistasis in the *Paramecium-Chlorella* symbiosis and test if compensatory evolution can rescue initially low fitness partner-switched symbioses. Using partner-switch experiments coupled with metabolomics we show evidence for intergenomic epistasis wherein low fitness arose from mismatched photoprotection traits and the resulting light stress experienced by non-native symbionts when in high light environments. Experimental evolution under high light conditions revealed that an initially low fitness partner-switched non-native host-symbiont pairing rapidly adapted, gaining fitness equivalent to the native host symbiont pairing in less than 50 host generations. Compensatory evolution took two alternative routes: Either, hosts evolved higher symbiont loads to mitigate for their new algal symbiont’s poor performance, or the algal symbionts themselves evolved higher investment in photosynthesis and photoprotective traits to better mitigate light stress. These findings suggest that partner-switching combined with rapid compensatory evolution will enable the recovery and local adaptation of symbioses in response to changing environments.

**Significance statement:** Symbiosis enables the formation of new organisms through the merger of once independent species. Through symbiosis, species can acquire new functions, driving evolutionary innovation and underpinning important ecosystem processes. Symbioses that breakdown due to changing environmental conditions can reform by acquiring new symbionts in a process called partner-switching but may exhibit low fitness due to their lack of coadaptation. Using a microbial symbiosis between the single-celled eukaryote *Paramecium* and the green alga *Chlorella* we show that low fitness in partner-switched host-symbiont pairings arises from mismatched photoprotection traits. However, such low fitness partner-switched pairings can be rapidly rescued by adaptive evolution, regaining high fitness in less than 50 host generations. Partner-switching coupled with rapid compensatory evolution can enable symbioses to recover from breakdown.

## Introduction

Beneficial symbioses have an inherent potential for conflict between the symbiotic partners. This can drive the breakdown of symbiosis if environmental conditions change the net benefit of interacting or if the pursuit of individual fitness favours cheating (1). Both situations can select for partner-switching to recombine novel symbiotic partnerships (2). Partner-switching can provide access to novel symbiotic phenotypes to overcome maladaptation to the prevailing environmental context (3) or restore symbiont function following breakdown (4, 5). The generation of phenotypic novelty through partner-switching arises from intergenomic epistasis (6); that is, genetic variation for the outcome of symbiosis in the form of host genotype by symbiont genotype interactions (G^H^ × G^S^) for symbiotic traits or fitness. Furthermore, the fitness effects of symbiosis are mediated by the environmental context (7), causing host-genotype-by-symbiont-genotype-by-environment interactions (G^H^ × G^S^ × E). A consequence of G^H^ × G^S^ × E interactions is that there is unlikely to be an optimal host-symbiont pairing across all environments, further driving selection for partner-switching or dynamic coevolution of the symbiosis (8). As such, partner-switching can enable niche-expansion by hosts (9, 10) and provide a mechanism by which host local adaptation can arise faster than through adaptation of the current symbiont (11, 12).

Newly interacting partner-switched symbioses are, however, unlikely to be co-adapted due to their lack of recent coevolutionary history and may, therefore, initially have low fitness. Indeed, despite the adaptive potential of partner-switching, new host-symbiont pairings, like genetic mutations, may more often be deleterious than beneficial to host fitness due to phenotypic mismatches or genetic incompatibilities. This has been observed in a range of symbiotic interactions: for example, a newly acquired *Symbiodinium* endosymbiont was found to translocate less fixed carbon than the native symbiont to its cnidarian host (13); novel bacterial endosymbionts had reduced vertical transmission rates in aphid hosts (14); and novel *Wolbachia* endosymbionts reduced the reproductive fitness of *Drosophila simulans* (15). How then do newly-formed, poorly co-adapted host-symbiont pairings become stable, beneficial symbioses? We hypothesise that rapid compensatory evolution could allow partner-switched symbioses to overcome their initially low fitness. Indeed, there is some, albeit limited, experimental evidence to support this idea: For example, the high fitness cost of newly acquired *Spiroplasma* endosymbionts in *Drosophila melanogaster* was ameliorated within only 17 host generations (16), although the underlying mechanisms of this fitness recovery remain unknown.

The microbial symbiosis between *Paramecium bursaria* and *Chlorella* provides an experimentally tractable model system to study intergenomic epistasis and the underlying molecular mechanisms. The ciliate host, *P. bursaria*, is a single-celled eukaryote, and each host cell contains 100-600 cells of the algal endosymbiont, *Chlorella* (17, 18). The *P. bursaria - Chlorella* symbiosis is based on a primary nutrient exchange of fixed carbon from the photosynthetic alga for organic nitrogen from the heterotrophic host (17, 19). This symbiosis is geographically widespread and genetically diverse, in part due to its multiple independent evolutionary origins (20, 21). The primary nutrient exchange is convergent among these origins, facilitating partner-switching, with concurrent divergence in other metabolic traits, causing phenotypic mismatches in partner-switched host-symbiont pairings (22). Here, using experimental partner-switches, we examined the pattern and mechanisms of intergenomic epistasis for three diverse host-symbiont strains, observing significant G^H^ × G^S^ × E interactions for host-symbiont growth rate and symbiont load, together with corresponding differences in metabolism. We then experimentally evolved a low fitness partner-switched host-symbiont pairing for ~50 host generations. We observed rapid compensatory evolution that improved fitness to equal to that of the native host-symbiont pairing mediated by evolved changes in symbiont load and metabolism.

## Results and Discussion

### Intergenomic epistasis for host-symbiont growth and symbiont load

We constructed all possible host-symbiont genotype pairings (n = 9) of 3 diverse strains of *Paramecium-Chlorella* and confirmed their identity by diagnostic PCR (Figure S1). We measured the growth reaction norm of each host-symbiont pairing across a light gradient (Figure 1a). All host-symbiont pairings showed the classic photosymbiotic reaction norm (23), such that growth rate increased with irradiance, but we observed significant intergenomic epistasis for host-symbiont growth rate (G^H^ × G^S^ × E interaction, ANOVA, F_17,162_ = 18.81, P<0.001). This was driven by contrasting effects of symbiont genotype on growth in the different host backgrounds across light environments. In the HK1 and HA1 host-backgrounds, similar growth reaction norms with light were observed for each symbiont genotype, whereas in the 186b host background the growth reaction norm varied according to symbiont genotype. Interestingly, the native 186b host-symbiont pairing had both the lowest intercept and the highest slope, indicating that in the 186b host background the native algal symbiont genotype was costlier in the dark yet more beneficial in high-light environments than non-native algal symbiont-genotypes.

**Figure 1.**
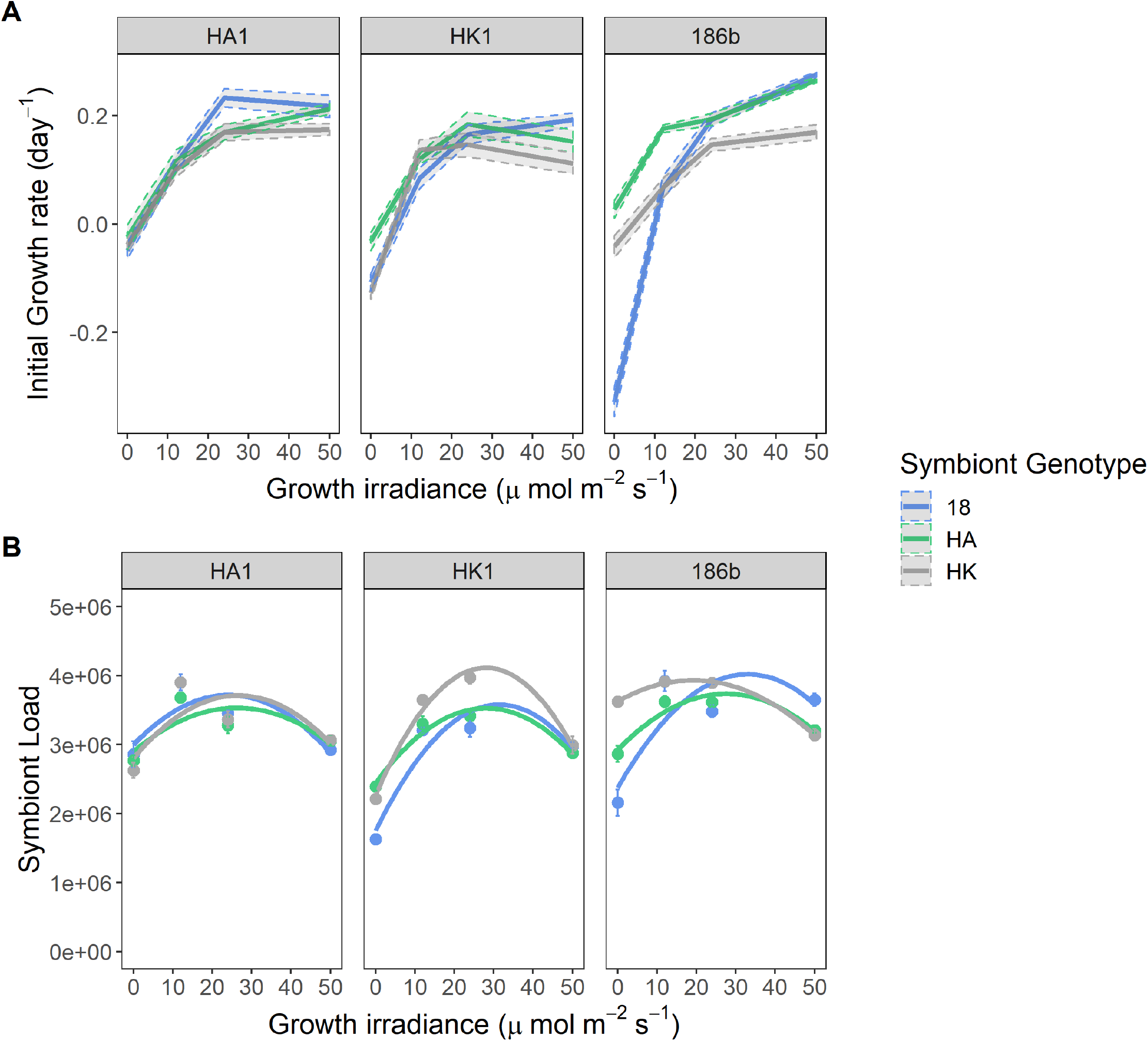
Growth rates and symbiont loads of the host-symbiont pairings. For both A and B, each panel presents the data for a specific genotype of *P. bursaria* host and the symbiont genotypes are distinguished by colour. A) Initial growth rates of the host-symbiont pairings across a light gradient. The data points show the mean (n=3) initial growth rate ±SE. B) Symbiont load of the host-symbiont pairings across a light gradient. The data points show the mean (n=3) symbiont load, measured as relative chlorophyll fluorescence, ±SE. The lines show polynomial models; the model coefficients showed a significant G^H^ × G^S^ interaction (ANOVA, F_8,36_ =27.22 (the intercept); 8.58 (first coefficient); 6.09 (second coefficient), P<0.001). For full statistical output see Table S1.

*P. bursaria* host cells regulate their algal symbiont load according to light irradiance to maximise the benefit-to-cost ratio of symbiosis, such that, for naturally occurring host-symbiont pairings, symbiont load peaks at low irradiance and is reduced both in the dark and at high irradiance (23–25). To test if regulation of symbiont load varied among host-symbiont pairings, we measured symbiont load across the light gradient as the intensity of single-cell fluorescence by flow cytometry (Figure 1b). All host-symbiont pairings showed the expected unimodal symbiont load curve with light, but nevertheless we observed significant intergenomic epistasis for symbiont load (G^H^ × G^S^ × E interaction, ANOVA, F_17,162_ = 3.78, P<0.001). Whereas, in the HA1 host similar symbiont load reaction norms were observed for each symbiont genotype, for the HK1 and 186b host backgrounds the form of the symbiont load reaction norms varied according to symbiont genotype. In the HK1 host, the magnitude of the symbiont load varied by symbiont genotype, such that higher symbiont loads were observed for the native compared to the non-native symbiont-genotypes. In the 186b host, peak symbiont load occurred at different light levels according to symbiont genotype, such that for the native symbiont the symbiont load curve peaked at a higher light intensity when compared to the non-native symbionts. (For the full output of the polynomial model, see Table S1.) This suggests that the HK1 and 186b host-genotypes discriminated among symbiont-genotypes, and then regulated symbiont load accordingly.

### Metabolic mechanisms of intergenomic epistasis

To investigate the potential metabolic mechanisms underlying the observed intergenomic epistasis we performed untargeted global metabolomics with ESI-ToF-MS independently for the host and symbiont metabolite fractions for each host-symbiont pairing across the light gradient (22). Light irradiance was the primary driver of differential metabolism for both host and symbiont, however, host-dependent differences in the metabolism of symbiont-genotypes could be detected. For the symbiont metabolite fraction subset by host-genotype, we observed native versus non-native clustering of symbiont metabolism only when associated with the 186b host-genotype (PCA Figures S2, OPLS-DA Figure S3). This is consistent with the larger phenotypic differences in growth and symbiont load observed among host-symbiont pairings with the 186b host-genotype compared to with either the HK1 or HA1 host-genotypes. Pairwise contrasts of the symbiont-genotypes in the 186b host-genotype background revealed a range of candidate symbiont metabolites which distinguished the native pairing from either non-native host-symbiont pairing. Putative identifications included, in the dark, elevated levels of candidate metabolites associated with stress responses (stress-associated hormones, jasmonic acid and abscisic acid, and stress associated-fatty acids, such as arachidonic acid) but reduced production of vitamins and co-factors by the native symbiont, compared to the non-native symbionts (Table S2). At high irradiance, the native symbiont showed higher levels of candidate metabolites in central metabolism, hydrocarbon metabolism and of biotin (vitamin B7), compared to the non-native symbionts (Table S3). In contrast, the non-native symbionts produced elevated levels, relative to native symbionts, of a candidate glutathione derivative; glutathione is an antioxidant involved in the ascorbate-glutathione cycle that combats high UV stress through radical oxygen scavenging (26, 27). Together, these data suggest that intergenomic epistasis can derive from mismatches in photoprotection and consequent responses to light stress by symbionts in novel host-symbiont pairings.

### Rapid compensatory evolution can rescue an initially low fitness partner-switched symbiosis

The partner-switched pairing of the 186b host with the HK1 symbiont showed substantially reduced growth at high light relative to the native 186b host-symbiont pairing. To test if this fitness deficit could be overcome through compensatory evolution, we established six replicate populations of each of these two symbiotic partnerships, which were propagated by weekly serial transfer for 25 transfers (approximately 50 host generations) at a high light regime (50μE; 14:10 L:D). The growth rate per transfer was higher for the native pairing than the non-native pairing (Figure S4) (linear mixed effect model, HK1 symbiont fixed effect of −0.08 ±0.006, T-value = −14.126, see Table S1 for full statistical output), but increased over time for both pairings (transfer number fixed effect 0.001 ±0.0004, T-value = 3.088). To test for adaptation, we compared the fitness effect of symbiosis at the beginning and the end of the transfer experiment by direct competition of either the ancestral or evolved host-symbiont pairings against the symbiont-free ancestral 186b host genotype across a light gradient. Starting fitness of symbiotic relative to non-symbiotic hosts increased more steeply with irradiance for the native than the partner-switched non-native pairing (Figure 2), but this difference had disappeared after evolution, such that both the native and non-native host-symbiont pairings showed increasing fitness relative to non-symbiotic hosts with increasing irradiance (symbiont genotype by light intensity by transfer number interaction term: ANOVA, F_7,45_ =6.20, P<0.001). Indeed, at 50 μE m^−2^ s^−1^, the light level used in the selection experiment, the large fitness deficit observed between the native and non-native pairing at the beginning of the experiment had been completely compensated. Comparison of the growth reaction norms of the evolving populations over time suggested that this amelioration occurred rapidly: By the tenth transfer, the native and non-native host-symbiont pairings showed equivalent growth responses to light (Welch t-test t(45.96) = −0.26, p = 0.80), in contrast to their substantially different ancestral growth reaction norms observed at the start of the evolution experiment (Welch t-test t(35.79) = 3.59, p = <0.001) (Figure S5). These data suggest that newly established partner-switched symbioses can rapidly achieve equivalent growth performance and fitness benefits as the native host-symbiont pairing by compensatory evolution.

**Figure 2.**
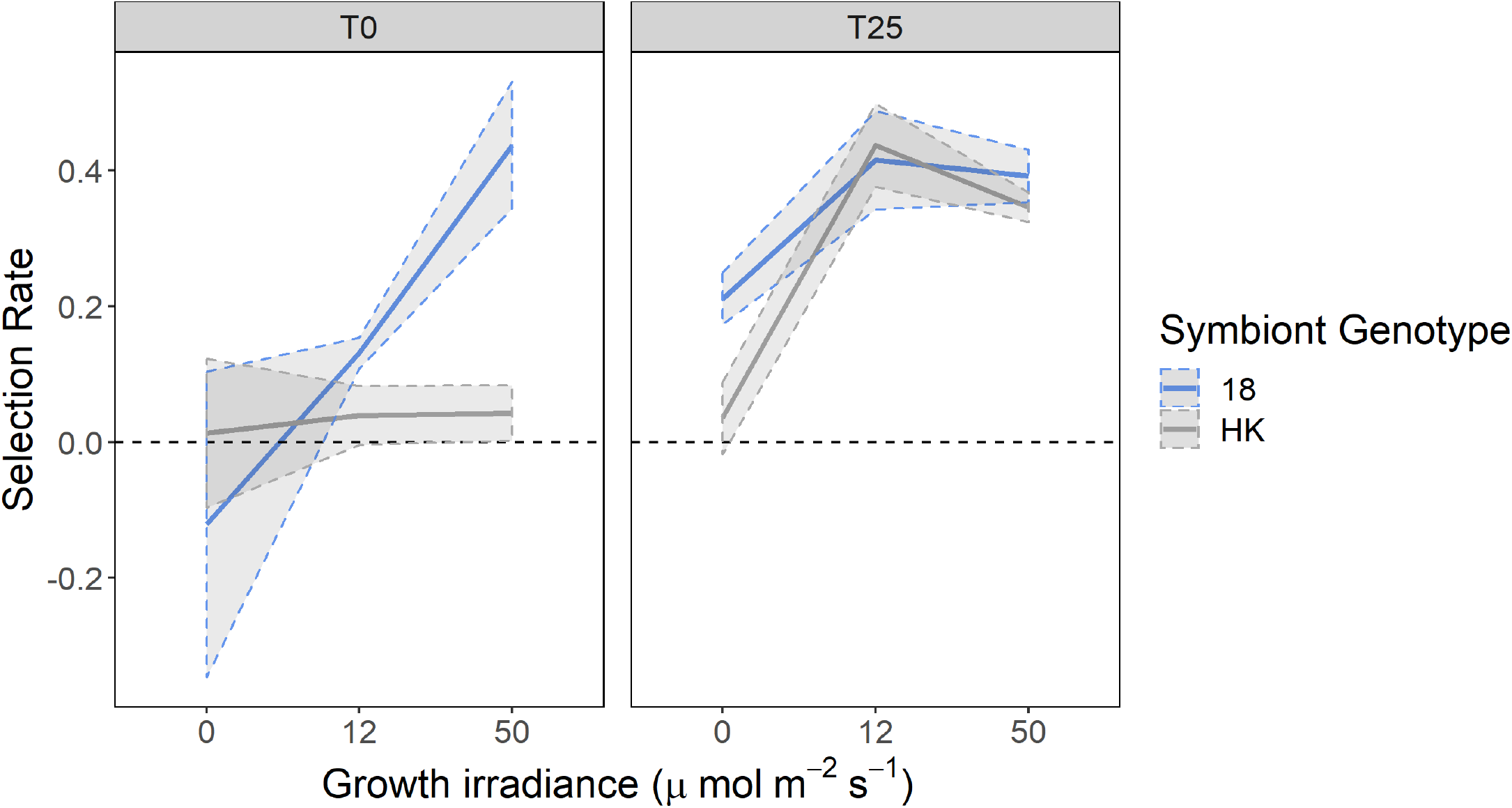
Fitness of the host-symbiont pairings relative to the symbiont-free host at the start and end of the evolution experiment. Lines show mean (n=6) competitive fitness of symbiont-containing hosts relative to the symbiont free 186b host calculated as selection rate, the shaded area denotes ± SE. The left-hand panel shows data measured at the start of the evolution experiment and the right-hand panel shows data measured at the end. Symbiont-genotype is denoted by colour. A selection rate above 0 indicates greater fitness in comparison to the symbiont-free host.

### Evolved changes in symbiont load regulation and metabolism

To understand the mechanisms of compensatory evolution, we first compared the symbiont load reaction norms of the ancestral and evolved native and non-native pairings (Figure 3). Both ancestral host-symbiont pairings showed the expected unimodal symbiont load curve with light, albeit with higher symbiont loads for the native compared to the non-native pairing at the highest light level, 50 μE m^−2^ s^−1^ irradiance, as used in the transfer experiment. Following evolution, the functional forms of the symbiont load reaction norms were altered in both the native and non-native pairings. Most notably, at 50 μE m^−2^ s^−1^ irradiance, whereas the non-native pairing had increased symbiont load, symbiont load had decreased in the native pairing, such that symbiont load was now higher in the non-native pairing (transfer by symbiont genotype interaction at high light: ANOVA, F_3,20_ = 16.88, P<0.001). Higher symbiont loads may therefore have contributed to the observed increased fitness of evolved compared to ancestral non-native pairings in the high light environment.

**Figure 3.**
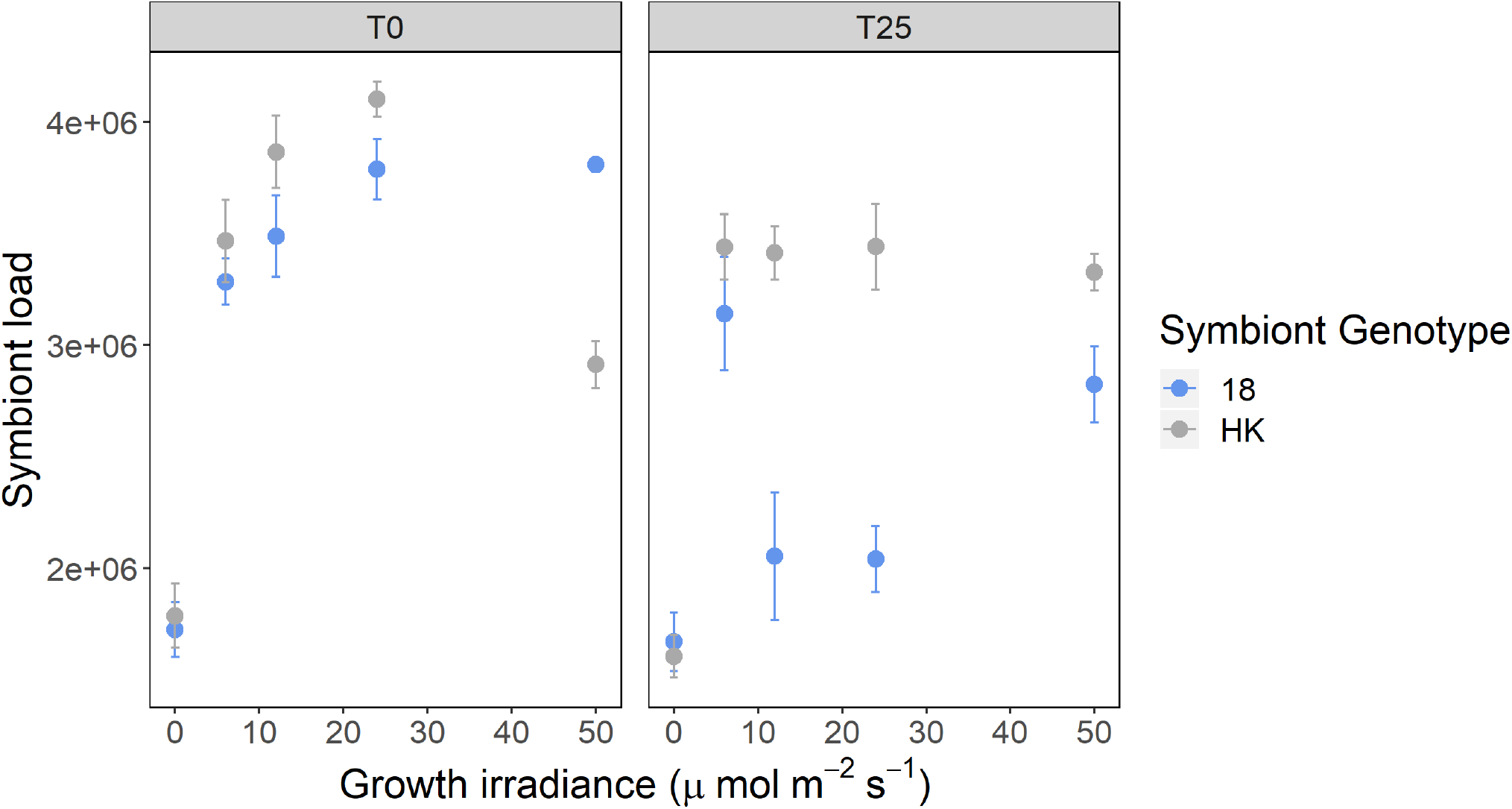
Symbiont load at the start and end of the evolution experiment. Symbiont load was measured across a light gradient. The left-hand panel shows data measured at the start of the evolution experiment and the right-hand panel shows data measured at the end. The points show the mean (n=6) relative chlorophyll fluorescence ± SE and symbiont genotype is denoted by colour.

Next, to investigate the underlying metabolic mechanisms, we performed untargeted metabolomics analyses on the separated *Chlorella* and *P. bursaria* fractions from samples taken the start and end of the evolution experiment grown at 50 μE m^−2^ s^−1^. The ancestral *P. bursaria* and *Chlorella* metabolic profiles of native and non-native host-symbiont pairings could be clearly distinguished. Following evolution, *P. bursaria* metabolism displayed a high degree of convergence between hosts evolved with the native versus the non-native symbionts (Figure 4a,c). This was driven by decreased levels of compounds of central metabolism (such as pyruvate and TCA cycle intermediates, antioxidants, lipids, and some amino acids) (Table S4), suggesting either increased pathway completion or a reduced metabolic rate, both of which can lead to increased efficiency. In addition, we observed increased levels of the amino acid cysteine and a shikimate pathway component in hosts evolved with the native versus the non-native symbionts (Figure S6). Levels of algal-cell degradation components (Figure S6), such as cell-wall degradation product chitotriose, were increased in some replicates of hosts evolved with either symbiont, potentially suggesting increased digestion of *Chlorella*, which is a known mechanism by which hosts control their symbiont load (28, 29).

**Figure 4.**
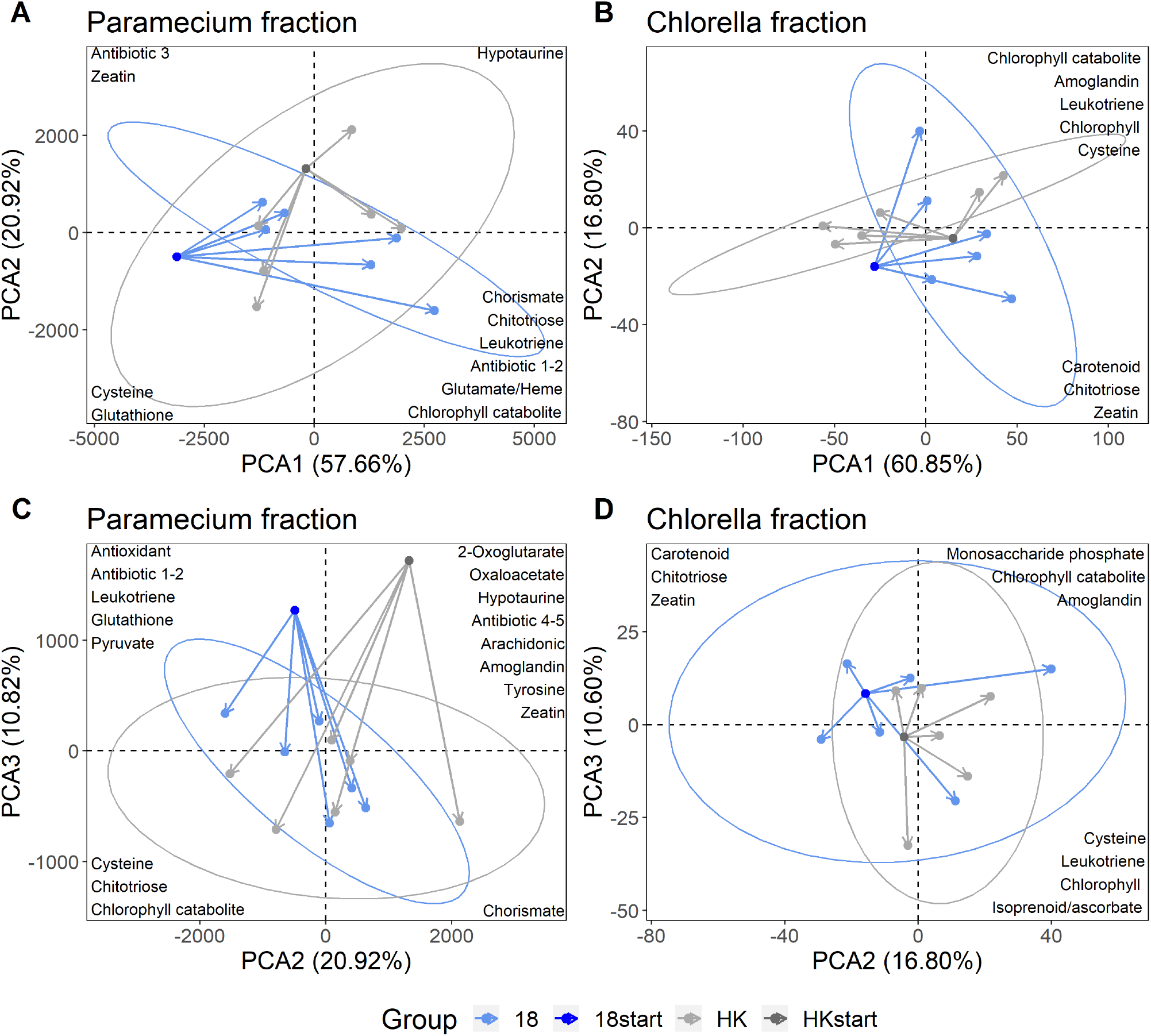
The trajectories of the metabolic profiles from the start to the end of the evolution experiment. These trajectories are shown within PCA plots and the arrows represent the movement in principal component space over the course of the experiment, with 95% confidence ellipses drawn for the evolved profiles. The metabolite identifications for the top loadings are shown in their corresponding location. Colour denotes the symbiont-genotype and shade represents whether the samples are from the start or end of the experiment. A and C show the results for the *P. bursaria* fraction, B and D the *Chlorella* fraction. The top row (A and B) plot PCA 1 versus PCA 2. The bottom row (C and D) plot PCA 2 versus PCA 3. The data here presents the biological replicates, which have been averaged over their technical replicates.

In contrast, evolved changes to the metabolic profiles of the algal symbiont genotypes showed less consistent differences among treatments (Figure 4b,d). Whereas all replicates of the native 186b *Chlorella* evolved in a similar direction, the replicates of the non-native HK1 *Chlorella* evolved in two different directions. Two of the HK1 replicates took a similar trajectory to the 186b symbionts, while the remaining four replicates all followed an alternative evolutionary trajectory. The group of four HK1 replicates that diverged during the experiment had lower production of metabolites within core aspects of metabolism, such as lipids, amino acids and carbohydrates. The second group, including the remaining two HK1 replicates and all the 186b replicates, had higher production of metabolites within primary metabolism pathways, particularly within lipids and carbohydrates, as well as a key chlorophyll compound, a photo-protective carotenoid, and secondary metabolites with potential antioxidant properties (Figure S7, Table S5). This greater investment into photosynthesis and photo-protection may improve carbon transfer to the host (30, 31), and decrease light stress, which aligns with the decrease in host antioxidants. Interestingly, the two HK1 replicates that converged metabolically with the native symbionts had a lower increase in symbiont load compared to the replicates that metabolically diverged (Table S6). This implies that the evolution of metabolism and symbiont load were linked, and that overall two alternative strategies of compensatory evolution emerged: either to have fewer, more beneficial symbionts or to have more, less-beneficial symbionts.

## Conclusion

Partner switching plays an important role in the evolution of a wide range of symbioses (2, 4, 5, 32, 33) enabling adaptation to changing environments and recovery from the breakdown of symbiosis. Because of intergenomic epistasis, partner-switched symbioses may possess novel adaptive phenotypes, but will sometimes exhibit low fitness due to mismatches between host and symbiont traits, owing to their lack of recent coevolutionary history (14, 15, 34). In the *Paramecium-Chlorella* symbiosis, low fitness following partner switching arose from mismatching photoprotection traits and the resulting light stress experienced by non-native symbionts when in high light environments, resulting in poor host-symbiont growth. This corresponds with findings from other photosynthetic symbioses, including coral-*Symbiodinium* and *Hydra-Chlorella*, where mismatching thermal and light stress tolerances contribute to the breakdown of symbiosis (35–38). Low fitness, partner-switched host-symbiont pairings were rescued by compensatory evolution, which took one of two routes: Either, hosts evolved higher symbiont loads to mitigate for their new algal symbiont’s poor performance, or the algal symbionts themselves evolved higher investment in photosynthesis and photoprotection traits to better mitigate light stress. Both strategies increased host-symbiont growth of the non-native pairing, leading to higher fitness equivalent to that of the native pairing. Together, these data suggest that, partner-switching combined with rapid compensatory evolution is likely to contribute to the recovery and local adaptation of symbioses in response to changing environments.

## Materials and Methods

### Cultures & Strains

The three natural strains used were: 186b (CCAP 1660/18) obtained from the Culture Collection for Algae and Protozoa (Oban, Scotland), and HA1 and HK1 isolated in Japan and obtained from the Paramecium National Bio-Resource Project (Yamaguchi, Japan). *P. bursaria* stock cultures were maintained at 25°c under a 14:10 L:D cycle with 50 μE m^−2^ s^−1^ of light. The stocks were maintained by batch culture in bacterized Protozoan Pellet Media (PPM, Carolina Biological Supply), made to a concentration of 0.66 g L^−1^ with Volvic natural mineral water, and inoculated approximately 20 hours prior to use with *Serratia marscesens* from frozen glycerol stocks.

To isolate *Chlorella* from the symbiosis, symbiotic cultures were first washed and concentrated with a 11μm nylon mesh using sterile Volvic. The suspension was then ultrasonicated using a Fisherbrand™ Q500 Sonicator (Fisher Scientific, NH, USA), at a power setting of 20% for 10 seconds sonification to disrupt the host cells. The liquid was then spotted onto Bold Basal Media plates (BBM) (39), from which green colonies were streaked out and isolated over several weeks. Plate stocks were maintained by streaking out one colony to a fresh plate every 3/4 weeks.

Symbiont-free *P. bursaria* were made by treating symbiotic cultures with paraquat (10 μg mL^−1^) for 3 to 7 days in high light conditions (>50 μE m^−2^ s^−1^), until the host cells were visibly symbiont free. The cultures were then extensively washed with Volvic and closely monitored with microscopy to check that re-greening by *Chlorella* did not occur. Stock cultures of the symbiont-free cells were maintained by batch culture at 25°c under a 14:10 L:D cycle with 3 μE m^−2^ s^−1^ of light and were given fresh PPM weekly.

### Cross infection

Symbiont-free populations of the three *P. bursaria* strains were re-infected by adding a colony of *Chlorella* from the plate stocks derived from the appropriate strain. This was done with all three of the isolated *Chlorella* strains to construct all possible host-symbiont genotype pairings (n=9). The regreening process was followed by microscopy and took between 2-6 weeks. Over the process, cells were grown at the intermediate light level of 12 μE m^−2^ s^−1^ and were given bacterized PPM weekly.

### Diagnostic PCR

The correct algae genotype within the cross-infections was confirmed using diagnostic PCR. The *Chlorella* DNA was extracted by isolating the *Chlorella* and then using a standard 6% Chelex100 resin (Bio-Rad) extraction method. A nested PCR technique with overlapping, multiplex Chlorophyta specific primers were used as described by Hoshina et al. (40). Standard PCR reactions were performed using Go Taq Green Master Mix (Promega) and 0.5μmol L^−1^ of the primer. The thermocycler programme was set to: 94°c for 5min, 30 cycles of (94°c for 30sec, 55°c for 30sec, 72°c for 60sec), and 5 min at 72°c.

### Growth rate

Growth rates of the symbioses were measured across a light gradient. The cells were washed and concentrated with a 11μm nylon mesh using sterile Volvic and re-suspended in bacterized PPM. The cultures were then split and acclimated to their treatment light condition (0, 12, 24, & 50 μE m^−2^ s^−1^) for five days. The cultures were then re-suspended in bacterized PPM to a target cell density of 150 cell mL^−1^. Cell densities were measured at 0, 24, 48 and 72 hours by fixing 360μL of each cell culture, in triplicate, in 1% v/v glutaraldehyde in 96-well flat-bottomed micro-well plates. Images were taken with a plate reader (Tecan Spark 10M) and cell counts were made using an automated image analysis macro in ImageJ v1.50i (41).

### Symbiont load

The symbiont load was measured in cultures derived from the growth rate experiment so that the data could be integrated between the two measurements. Triplicate 300μl samples of each cell culture were taken from 72-hour cultures for flow cytometry analysis. Host symbiont load was estimated using a CytoFLEX S flow cytometer (Beckman Coulter Inc., CA, USA) by measuring the intensity of chlorophyll fluorescence for single *P. bursaria* cells (excitation 488nm, emission 690/50nm) and gating cell size using forward side scatter; a method established by Kadono et al. (18). The measurements were calibrated against 8-peak rainbow calibration particles (BioLegend), and then presented as relative fluorescence to reduce variation across sampling sessions.

### Partner-switching - Metabolomics

Cultures of the symbiotic pairings were washed and concentrated with a 11μm nylon mesh using sterile Volvic and re-suspended in bacterized PPM. The cultures were then split and acclimated at their treatment light condition (0, 12 & 50 μE m^−2^ s^−1^) for seven days. The symbiotic partners were separated in order to a get *P. bursaria* and *Chlorella* metabolic fraction. The *P. bursaria* cells were concentrated with a 11μm nylon mesh using Volvic and then the *P. bursaria* cells were disrupted by sonication (20% power for 10 secs). 1ml of the lysate was pushed through a 1.6μm filter, which caught the intact *Chlorella* cells, and the run-through was collected and stored as the *P. bursaria* fraction. The 1.6μm filter was washed with 5ml cold deionized water, and then reversed so that the *Chlorella* cells were resuspended in 1ml of cold methanol, which was stored as the *Chlorella* fraction. After which the *Chlorella* fraction samples were already in methanol, but the *P. bursaria* fraction samples had then to be diluted by 50% with methanol.

Metabolic profiles were recorded using ESI ToF-MS, on the Qstar Elite with automatic injection using Waters Alliance 2695 HPLC (no column used), in positive mode. This is an established high-throughput method with a large mass range (50 Da to 1000 Da).

Mass spectrometry settings:

|  |  |
| --- | --- |
| Polarity: | positive |
| Ion Spray voltage: | 4.2 kV |
| Declustering potential: | 120 V |
| Focusing potential: | 265 V |
| Source temperature: | 200°C |
| Gas Flow: | 40 ml min<sup>-1</sup> |
| Solvent: | 50:50 methanol to water at flow rate 40$\mu\text{l min}^{-1}$ |
| Injected volume: | 10$\mu\text{l}$ |

The processing was performed using in-house software Visual Basic macro 216 (42), which combined the spectra across the technical replicates by binning the crude m/z values into 0.2-unit bins. The relative mass abundances (% total ion count) for each bin was summed. Pareto scaling was applied to the results, and the data was then analysed by principal component analysis using SIMCA-P software (Umetrics). When treatment-based separation was observed, supervised orthogonal partial least squares discriminant analysis (OPLS-DA) separation was then performed using the discriminatory treatment with the SIMCA-P software.

### Partner-switching - Identification of significant masses

Masses of interest were annotated using the initial identifications from the in-house software program and further comparisons against KEGG (https://www.genome.jp/kegg/) (43, 44) and Metlin (https://metlin.scripps.edu) (45) databases. The Metabolomics Standards Initiative requires two independent measures to confirm identity, this partner-switching metabolomic analysis only used one measure (accurate mass) and therefore, meets only the level 2 requirements of putative annotated compounds.

### Evolution Experiment

The populations used derive from the cross-infections and, therefore, the symbiotic partnerships come from the same cured 186b ancestor that was then re-infected with either its native (186b) or novel (HK1) symbionts. The two symbiotic partnerships were split into six replicate populations that were used as the starting populations. The 200ml populations were propagated by weekly serial transfer for 25 transfers at a high light (50 μE m^−2^ s^−1^) 14:10 L:D cycle. At every transfer, cell-density was equalised to 100 cells mL^−1^ and the transferred cells were washed with a 11μm nylon mesh using Volvic before being re-suspended in bacterized PPM. Cell density was measured before and after each transfer by fixing 360μL of each cell culture, in triplicate, in 1% v/v glutaraldehyde in 96-well flat-bottomed micro-well plates. Images were taken with a plate reader (Tecan Spark 10M) and cell counts were made using an automated image analysis macro in ImageJ v1.50i (41). Growth rate and symbiont load assays were conducted at the start, T10, T20 and end of the experiment using the method described above.

### Evolution experiment - Fitness assay

Fitness assays were conducted at the start and end of the evolution experiment. *P. bursaria* cultures, both the symbiotic pairings and the symbiont-free ancestor, were washed with Volvic and resuspended in bacterized PPM. The cultures were then split and acclimated at their treatment light level (0,12,50 μE m^−2^ s^−1^) for five days. Cell densities were counted by fixing 360μL of each cell culture, in triplicate, in 1% v/v glutaraldehyde in 96-well flat-bottomed micro-well plates. Images were taken with a plate reader (Tecan Spark 10M) and cell counts were made using an automated image analysis macro in ImageJ v1.50i (41). The competitions were started by setting up microcosms that each contained 50:50 populations of green and white cells (with target values of 20 green cells and 20 white cells per mL) that were in direct competition. Cells were sampled on day 0 and day 7 on a flow cytometer and the proportion of green to white cells was measured and used to calculate the selection rate. Green versus white cells were distinguished using single cell fluorescence estimated using a CytoFLEX S flow cytometer (Beckman Coulter Inc., CA, USA) by measuring the intensity of chlorophyll Fluorescence (excitation 488nm, emission 690/50nm) and gating cell size using forward side scatter; a method established by Kadono et al. (18). The measurements were calibrated against 8-peak rainbow calibration particles (BioLegend), and then presented as relative fluorescence to reduce variation across sampling sessions. The re-establishment of endosymbiosis takes between 2-4 weeks, and this method was tested to ensure that the symbiont-free cells do not re-green over the course of the experiment.

### Evolution experiment - Metabolomics

The cultures were sampled at the start and end of the evolution experiment. Cultures were washed and concentrated with a 11μm nylon mesh using Volvic and re-suspended in bacterized PPM. The cultures were acclimated at their treatment light condition (50 μE m^−2^ s^−^ 1) for seven days. For the start point, the six experimental replicates were used as replicates for the metabolomics. For the end point, three replicates of each of the six experimental replicates were used for the metabolomics because divergence may have occurred over the course of the experiment. At each sampling event, the symbiotic partners were separated in order to a get *P. bursaria* and *Chlorella* metabolic fraction using the extraction method described above. Samples were freeze-dried for storage, and then resuspended in 50:50 methanol to water prior to mass spectrometry.

The samples were analysed with a Synapt G2-Si with Acuity UPLC, recording in positive mode over a large untargeted mass range (50 – 1000 Da). A 2.1×50mm Acuity UPLC BEH C18 column was used with acetylnitrile as the solvent. The machine settings are listed in detail below:

Mass spectrometry settings:

|  |  |
| --- | --- |
| Polarity: | positive |
| Capillary voltage: | 2.3 kV |
| Sample Cone voltage: | 20 V |
| Source Temperature: | 100°C |
| Desolvation temperature: | 280°C |
| Gas Flow: | 600 L hr<sup>-1</sup> |
| Injected volume: | 5µl |
| Column temperature: | 45°C |

Gradient information:

| Time (mins) | Water (%) | Acetonitrile (%) |
| --- | --- | --- |
| 0 | 95 | 5 |
| 3 | 65 | 35 |
| 6 | 0 | 100 |
| 7.5 | 0 | 100 |
| 7.6 | 95 | 5 |

The *P. bursaria* and *Chlorella* fraction were analysed separately. The xcms R package (46– 48) was used to extract the spectra from the CDF data files, using a step argument of 0.01 m/z. Peaks were identified, and then grouped across samples. These aligned peaks were used to identify and correct correlated drifts in retention time from run to run. Pareto scaling was applied to the resulting intensity matrix.

### Evolution experiment - Metabolomics analysis

The metabolic profiles from the start and end of the experiment were compared using principal component analysis (PCA) with the prcomp() function in Base R (https://www.rproject.org/). For both fractions the first three components were considered, this accounted for >88% of the variance. The top 1% of the loadings were selected using the absolute magnitude of the loadings. These top loadings were identified where possible, and the identified loadings were then depicted in their associated component space. The relative abundance of these top loadings was visualised using heatmaps drawn with the heatmap.2() function from the gplot package (49). The phylogenies were based on UPGMA clustering of the PCA coordinates of the samples using the hclust() function. This approach of integrating metabolic data and genotypes in heatmaps has been used previously (50).

### Evolution experiment - Identification of significant masses

Masses of interest were investigated using the MarVis-Suite 2.0 software (http://marvis.gobics.de/) (51), using retention time and mass to compare against KEGG (https://www.genome.jp/kegg/) (43, 44) and MetaCyc (https://biocyc.org/) (52) databases. The Metabolomics Standards Initiative requires two independent measures to confirm identity, which the combination of retention time and accurate mass achieves for the analysis of the evolution experiment metabolomics.

## Data Analysis

Statistical analyses were performed in Rv.3.5.0 (53) and all plots were produced using package ggplot2 (54) unless otherwise stated. Physiology tests were analysed by both ANOVA and ANCOVA, with transfer time, host and symbiont identity as factors. A linear mixed effect model was used to analysis the growth rate per transfer using lm() function from the nlme package (55). The lm model included fixed effects of symbiont genotype and transfer number, and random effects of transfer number given sample ID. Details of the statistical methods used are within the supplementary statistics table (Table S1).

## Acknowledgements

This work was funded by grants NE/K011774/2 and NE/V000128/1 from the Natural Environment Research Council, UK to M.A.B, D.D.C, and A.J.W and a White Rose DTP studentship from the Biotechnology and Biological Sciences Research Council, UK (BB/011151/1) to M.E.S.S. The funders had no role in the design of the study, the collection, analysis and interpretation of data or writing of the manuscript. We are grateful to Heather Walker for her technical assistance with the mass spectrometry.

