## Supplementary Information for "Rapid compensatory evolution can rescue low fitness symbioses following partner-switching"

**This PDF file includes:**

Figures S1 to S7

Tables S2 to S6

**Other supplementary materials for this manuscript include the following:**

Statistical results Table S1

**L	s18	sHA	sHK	h18	hHA	hHK	k18	kHA	kHK**

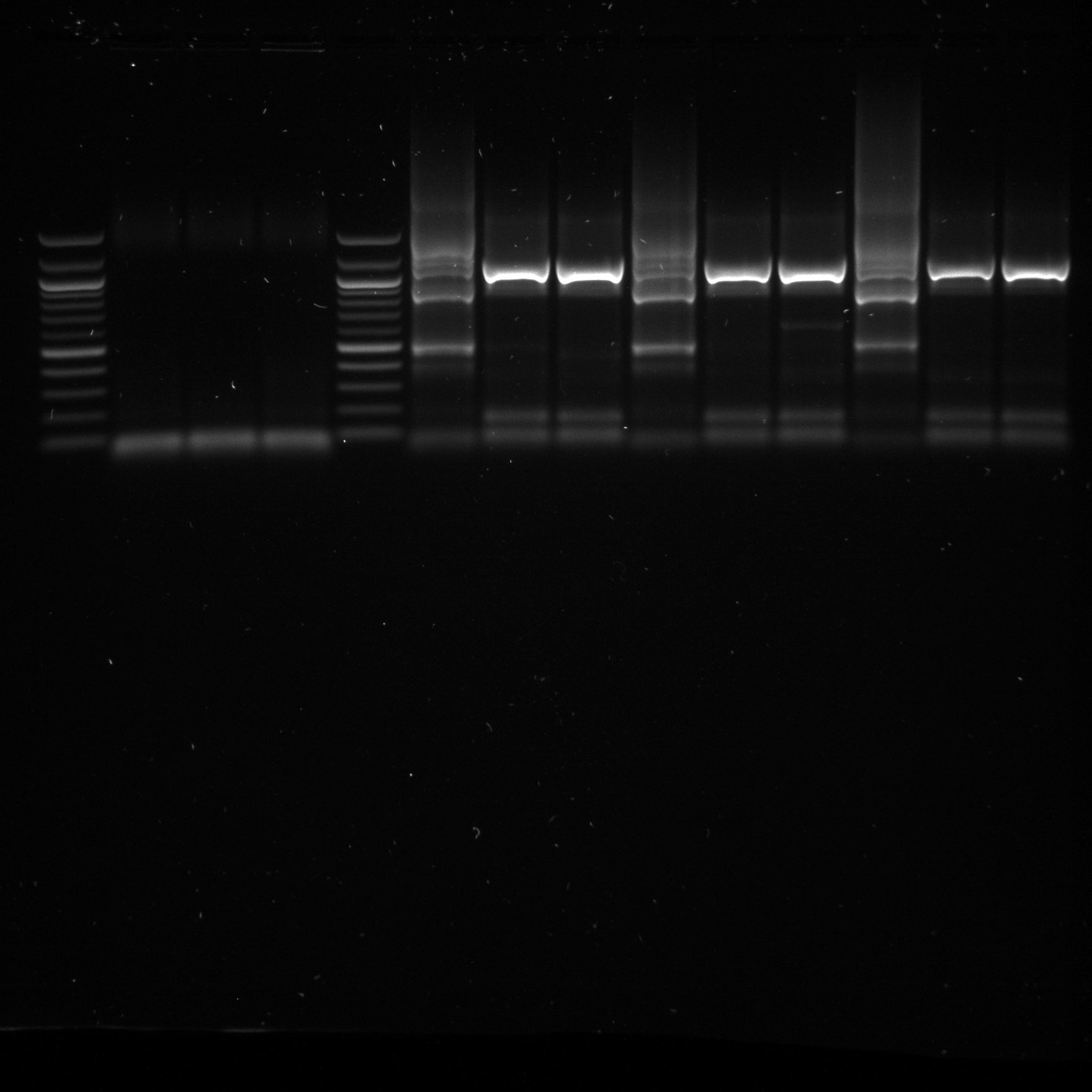

**186b host**

**HA1 host**

**HK1 host**

**1000 bp**

**500 bp**

**Figure S1. PCR confirmation of symbiont-genotype within the reciprocal cross infections.** Overlapping, multiplex primers were used to amplify fragments within the 18S rDNA and ITS region of the *Chlorella* nuclear genome. In this region the ‘American/Japanese’ strains, such as HA1 and HK1, have had three introns inserted that the ‘European’ strains, such as 186b, lack (Hoshina and Imamura, 2008; Hoshina et al., 2005). The banding pattern results here confirm that the cross-infections were successful and contain the correct *Chlorella* genotype, specifically that the distinct banding pattern of 186b was present when expected. This PCR method can distinguish between ‘American/Japanese’ and ‘European’ strains, but not between strains that come from the same biogeographical clade. Host genotype has been shortened to a letter (‘s’ = 186b host, ‘h’ = HA1 host, ‘k’ = HK1 host); symbiont genotype is shown by two capitals (‘18’ = 186b symbiont, ‘HA’ = HA1 symbiont, ‘HK’ = HK1 symbiont. Shown alongside a 100bp ladder.

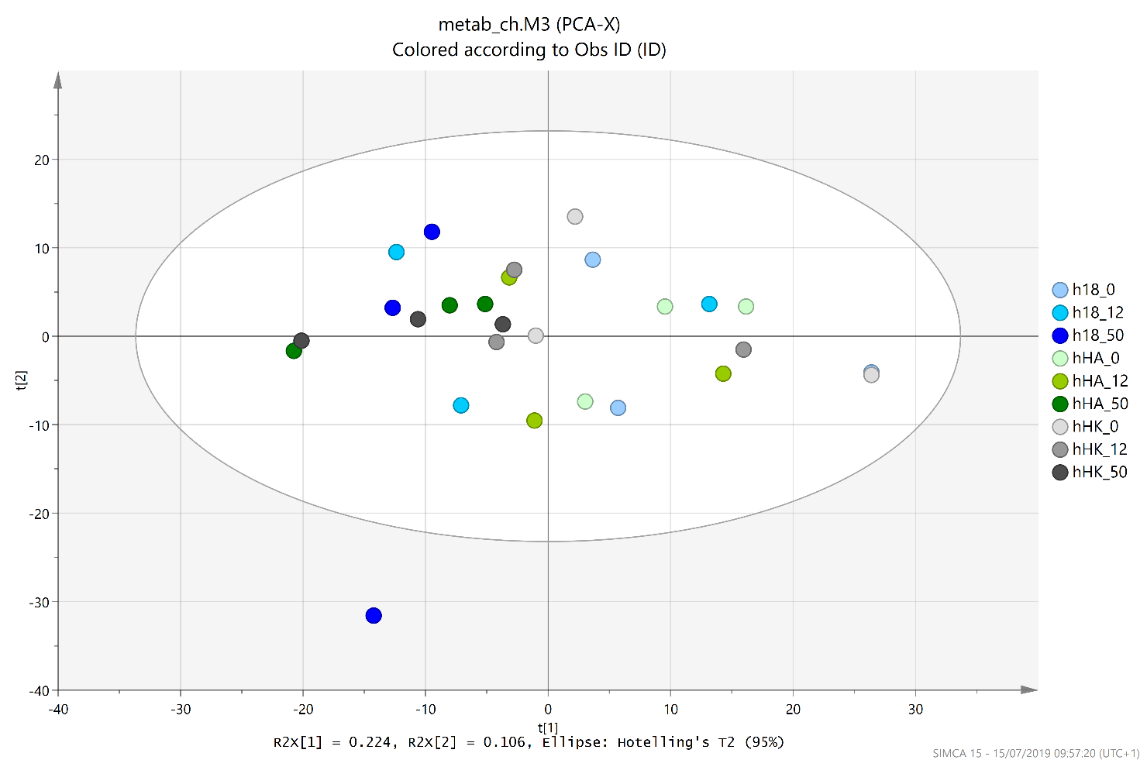

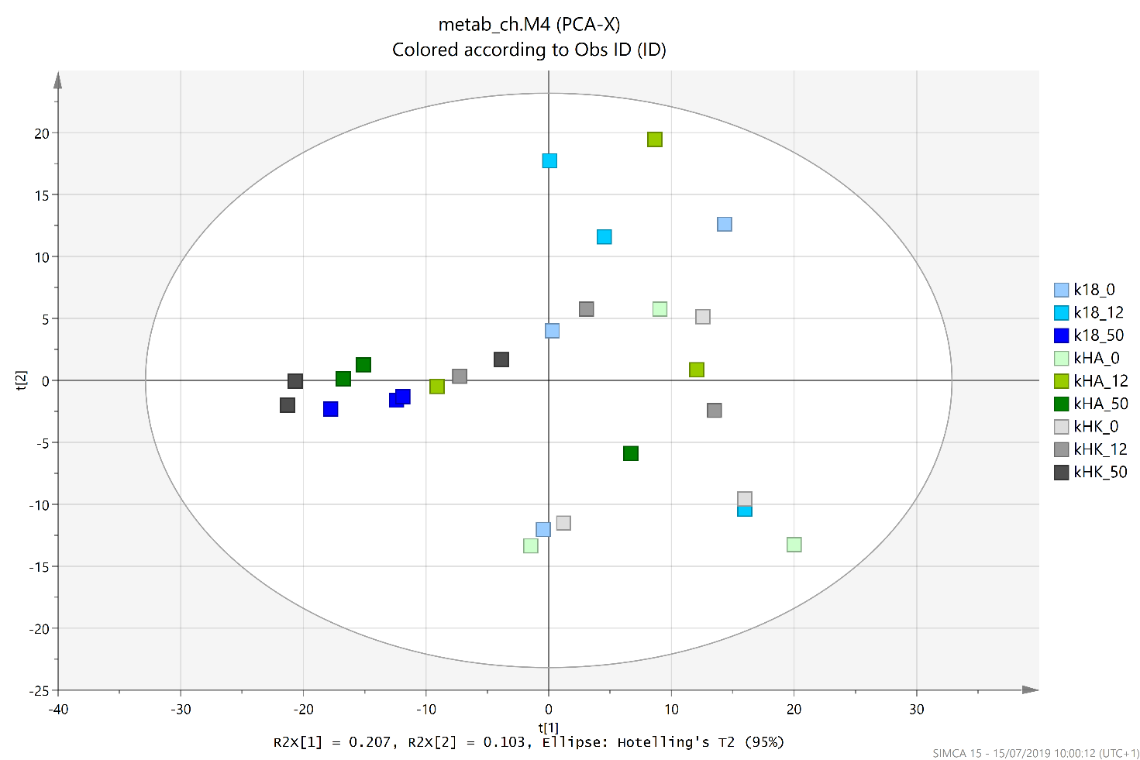

**A**

**B**

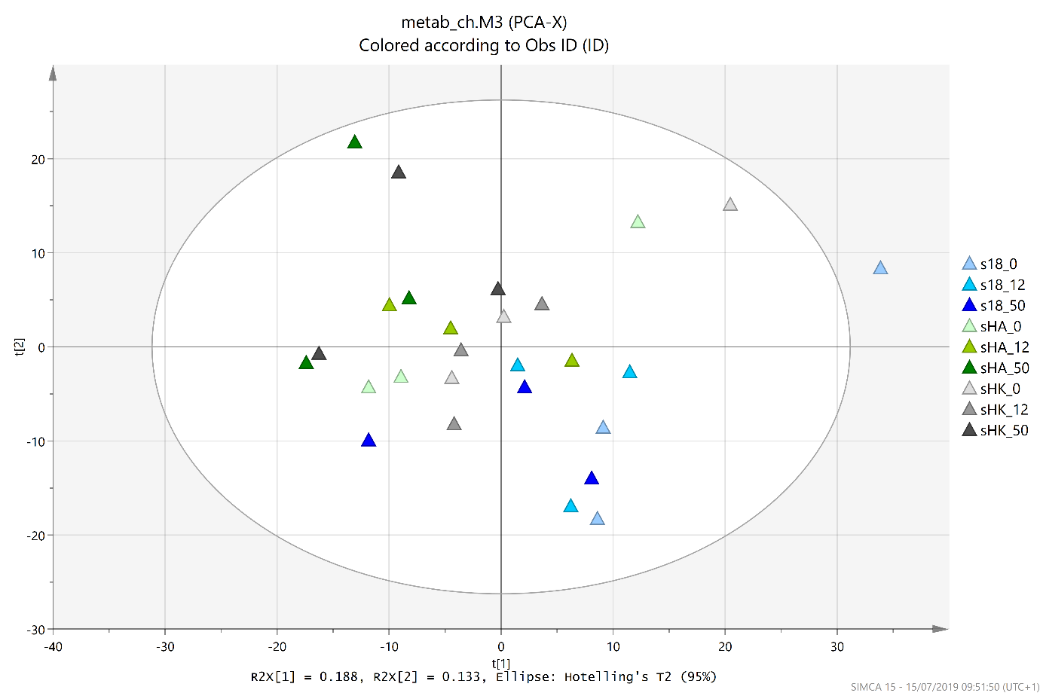

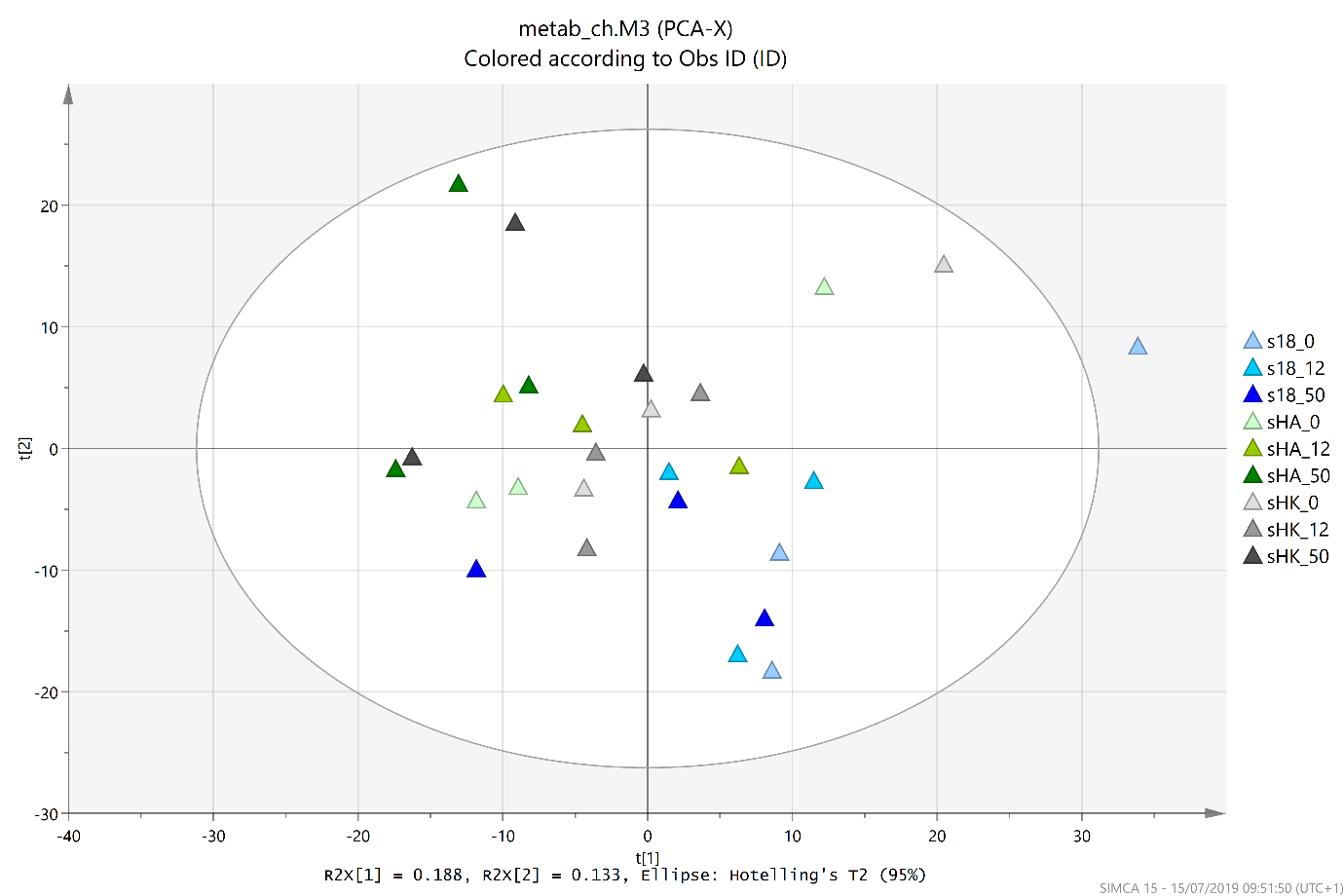

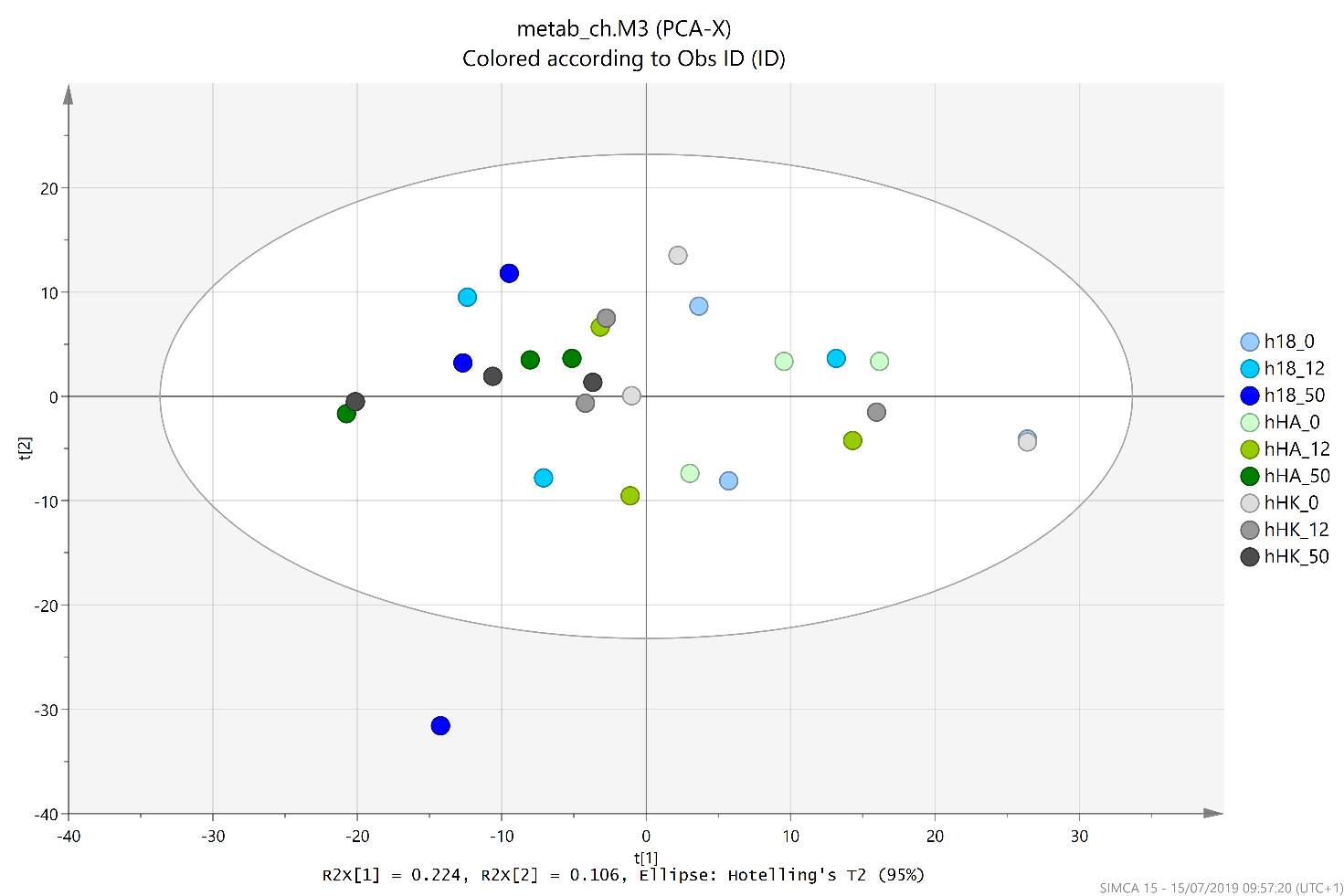

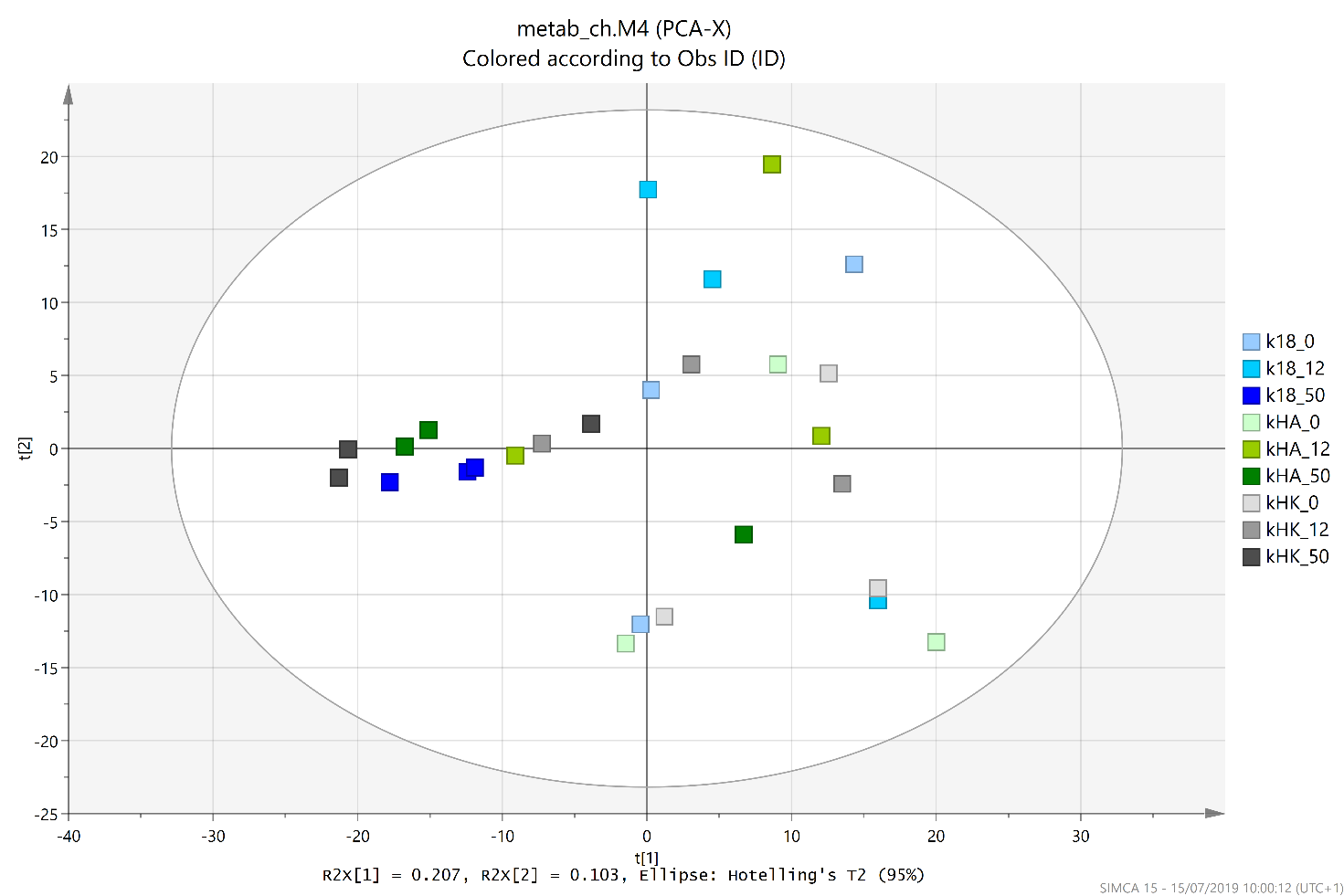

**C**

**Figure S2.** **Clustering patterns of the *Chlorella* metabolic fraction subset by host-genotype.** These PCA plots show the HA1 host (A), the HK1 host (B), and the 186b host (C). Each point represents the metabolic profile of a sample; with the shape denoting the *P. bursaria* host genotype, the colour denoting the *Chlorella* symbiont genotype and the colour shade denoting the light intensity. Only within the 186b host (C) do the samples clusters by colour, and therefore, symbiont genotype. There are 3 replicates of each combination of host, symbiont and light intensity.

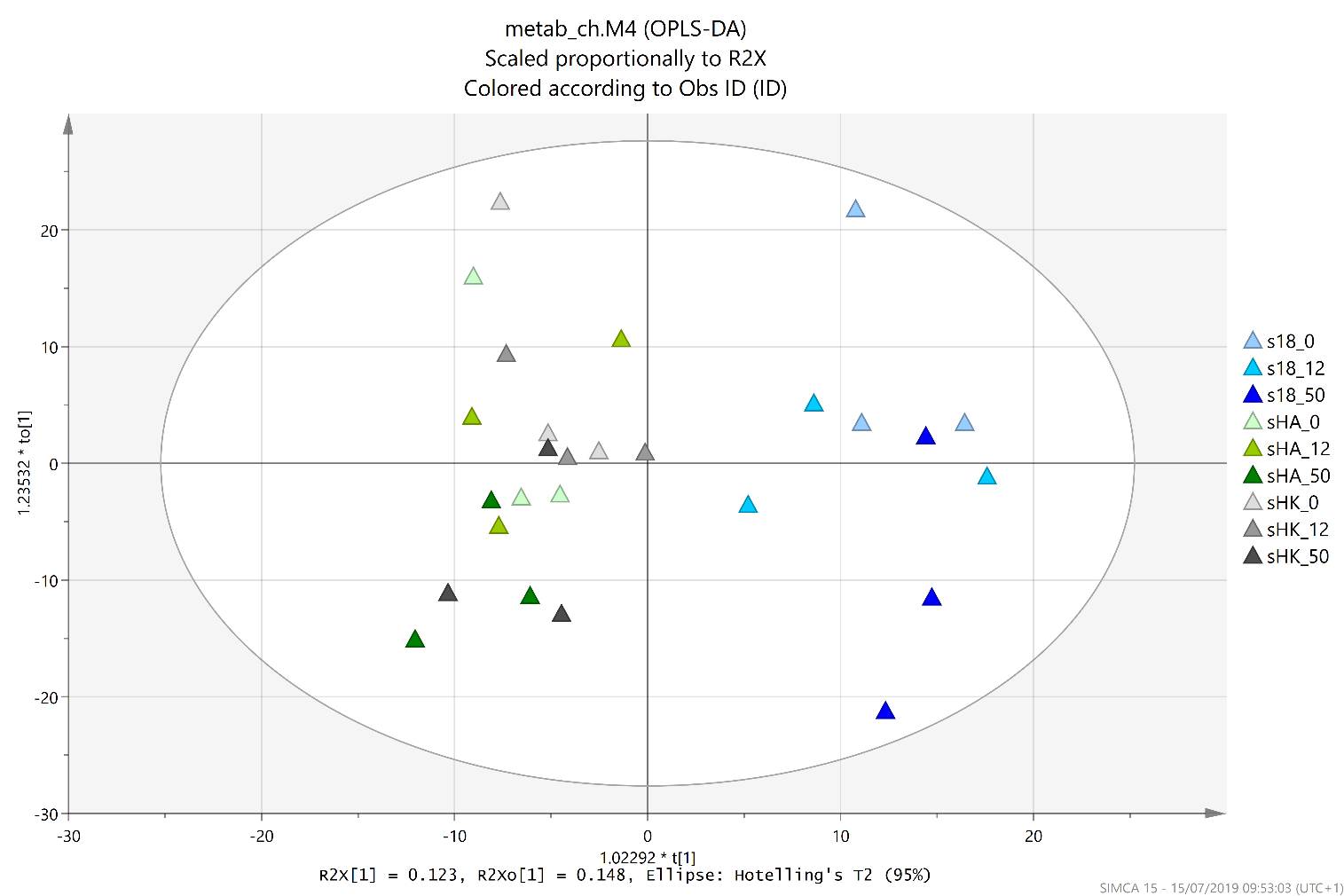

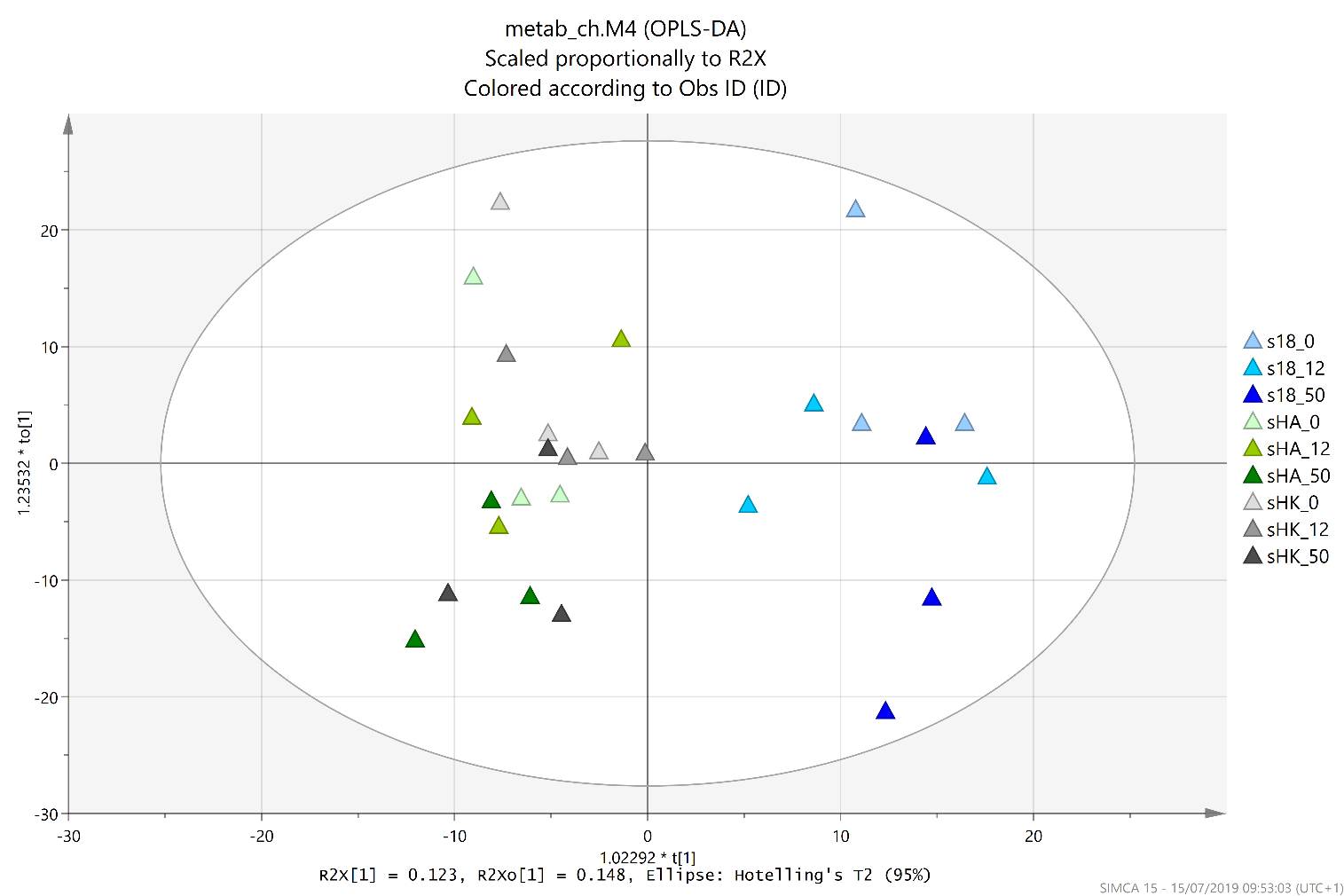

**Figure S3. Separation by symbiont-genotype within the 186b host subset of the *Chlorella* metabolic fraction.** This OPLS-DA plot follows the initial clustering by symbiont-genotype within the 186b host subset in the PCA plot (Figure S2 C). Each point represents the metabolic profile of a sample; with the colour denoting the *Chlorella* symbiont genotype and the shade of the colour denoting the light intensity. The samples separate between the ‘blue’ samples (186b symbiont-genotype) and the ‘green’ and ‘grey’ samples (HA1 and HK1 symbiont genotypes). There are 3 replicates of each combination of host, symbiont and light intensity.

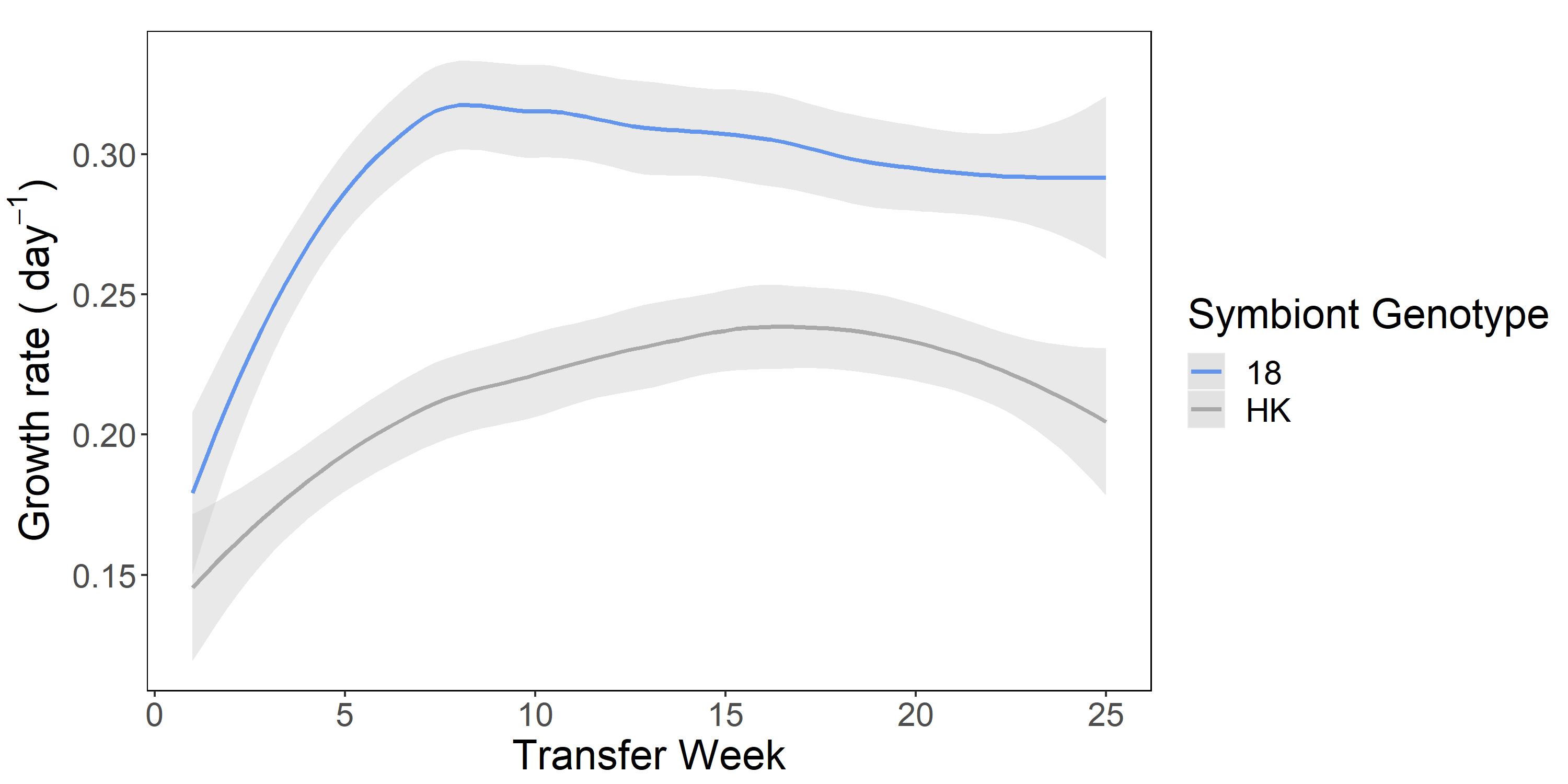

**Figure S4. Weekly growth rates of the native and novel symbioses across the evolution experiment.** The lines show the smoothed mean (n=6) growth rates ± SE and colour denotes the symbiont genotype (blue = 18 = native symbiont; grey = HK = novel symbiont). The smoothing function used was the loess method.

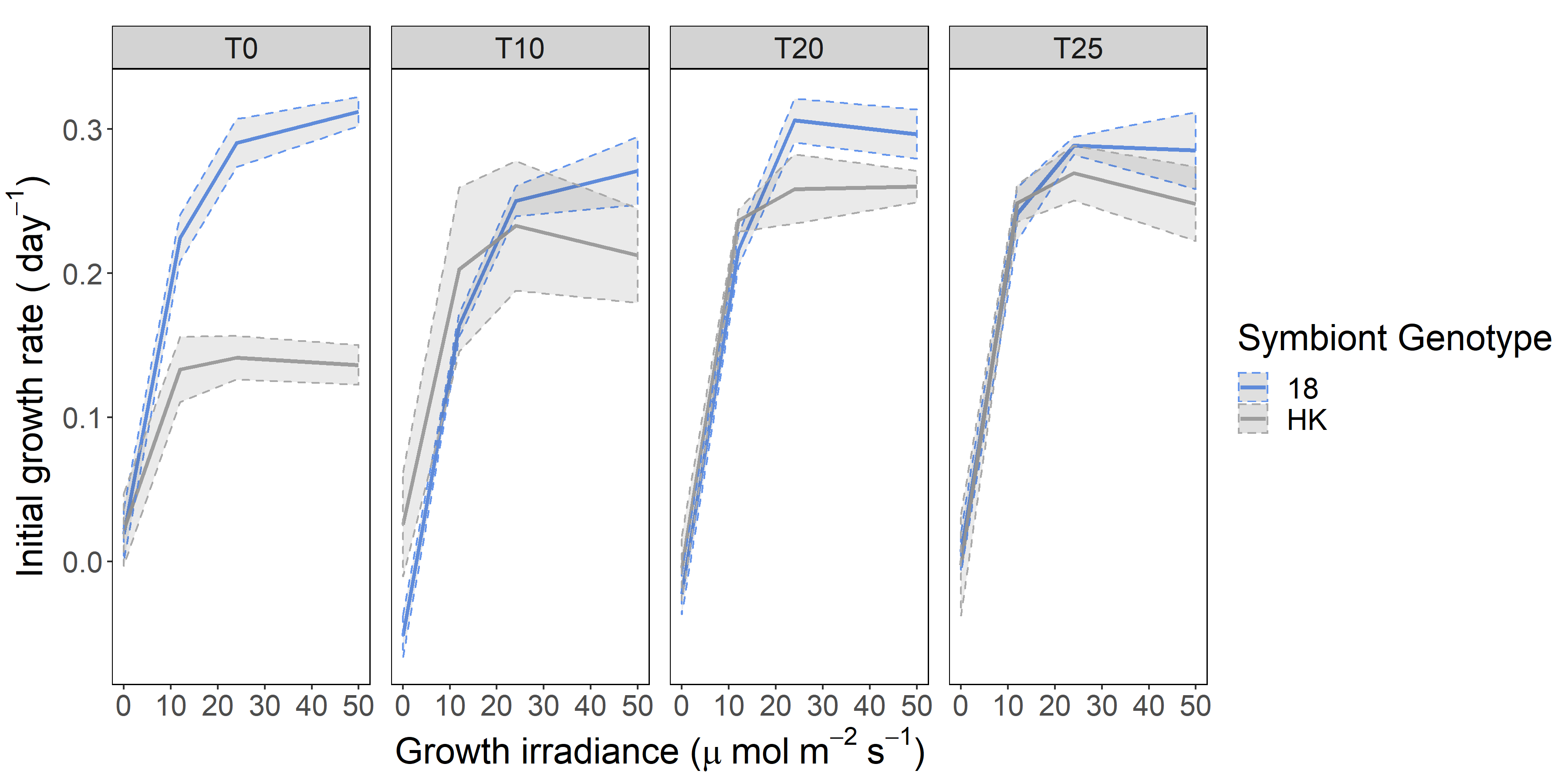

**Figure S5. Growth rate assays performed at multiple points throughout the evolution experiment.** Each panel shows the mean (n=6) initial growth rate across a light gradient and the shaded area denotes ± SE. The panels represent the transfer week within the evolution experiment at which the growth assay was performed (T0 = week 0, T10 = week 10, T20 = week 20 & T25 = week 25). The symbiont-genotype is denoted by colour.

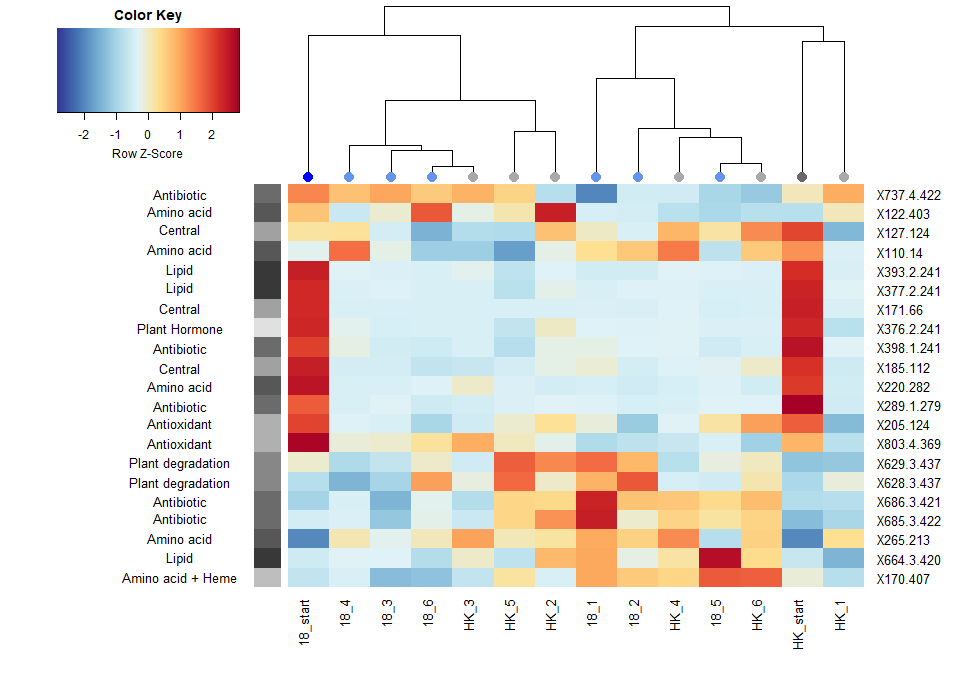

**Figure S6. Metabolites of interest across the start and end of the evolution**

**experiment within the *P. bursaria* fraction.** The data is shown as a heatmap with the colour representing the relative abundance of the metabolites. The metabolites depicted were identified from the top loadings of the PCA plots. The columns in the heatmap correspond to sample; these are labelled with their symbiont-genotype (18 = 186b; HK = HK1) and with the replicate number if from the end of the experiment or ‘start’ if from the start. The column on the left of the heatmap indicates the function of the metabolites and the column on the right indicates the loading ID, which corresponds to the identification Table S4. The phylogeny of the samples was calculated with their principal component coordinates using UPGMA clustering, and the order of the rows was assigned by UPGMA clustering performed on the rows’ distance measures (based on the Pearson correlation co-efficient).

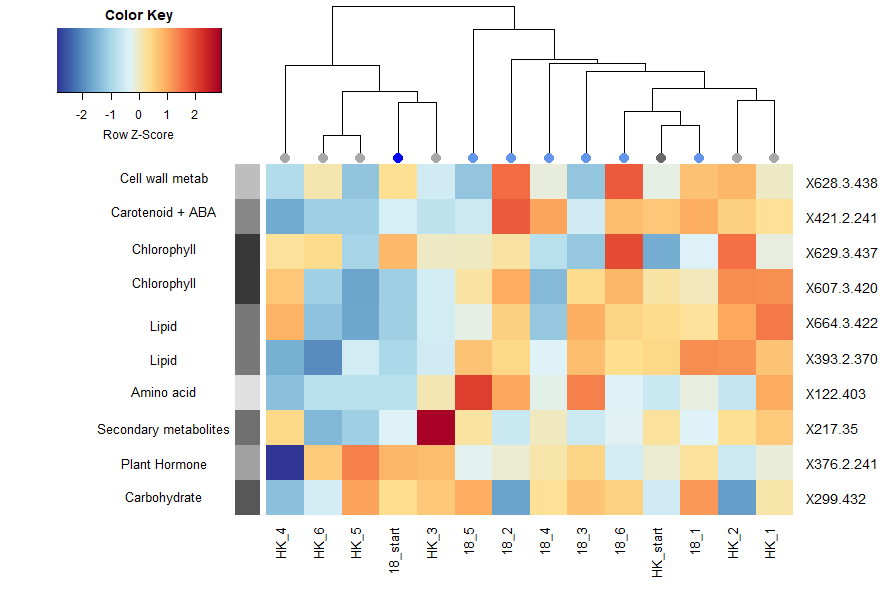

**Figure S7.** **Metabolites of interest across the start and end of the evolution**

**experiment within the *Chlorella* fraction.** The data is shown as a heatmap with the colour representing the relative abundance of the metabolites. The metabolites depicted were identified from the top loadings of the PCA plots. The columns in the heatmap correspond to the sample; these are labelled with their symbiont-genotype (18 = 186b; HK = HK1) and with the replicate number if from the end of the experiment or ‘start’ if from the start. The column on the left of the heatmap indicates the function of the metabolites and the column on the right indicates the loading ID, which corresponds to the identification Table S5. The phylogeny of the samples was calculated on their principal component coordinates using UPGMA clustering, and the order of rows was assigned by UPGMA clustering performed on the rows’ distance measures (based on the Pearson correlation co-efficient).

**Table S2** **Symbiont-genotype specific metabolites in the dark within the 186b *P. bursaria* host.** These metabolite IDs were highlighted by the pairwise contrasts and were found to have significantly higher abundances in one symbiont-genotype compared to another within the 186b host subset of the *Chlorella* metabolic fraction in the dark (0µE).

| **Strain Associated** | **Comparison** | **mz ID** | | **Detected Mass** | | **Accurate Mass** | | **Adduct** | | **Candidate Compound** | | **Pathway** | | **Stress Associated** | |
| --- | --- | --- | --- | --- | --- | --- | --- | --- | --- | --- | --- | --- | --- | --- | --- |
| **s18** | s18 vs sHA | 118 | | 117.966 | | 117.0426 | | H+ | | Aspartate-4-semialdehyde | | Amino acid | |  | |
|  |  |  | |  | | 117.0578 | | H+ | | Indole | | Amino acid + hormone | |  | |
|  |  |  | |  | | 117.0790 | | H+ | | Glycinebetaine | | Amino acid + osmolyte | |  | |
|  |  |  | |  | | 117.0790 | | H+ | | Valine | | Amino acid | |  | |
|  | s18 vs sHA | 134.2 | | 134.109 | | 133.1040 | | H+ | | Aspartate | | Amino acid | |  | |
|  | s18 vs sHA | 255.2 | | 255.104 | | 216.1725 | | K+ | | w-hydroxydodecanoic acid | | Hydroxy fatty acids | |  | |
|  |  |  | |  | | 254.2246 | | H+ | | Palmitoleic acid | | Unsaturated fatty acids | |  | |
|  | s18 vs sHA | 343.2 | | 343.153 | | 342.1162 | | H+ | | Disaccaride | | Carbohydrate | |  | |
|  |  |  | |  | | 304.2402 | | K+ | | Arachidonic acid | | Unsaturated fatty acids | | Yes | |
|  |  |  | |  | | 304.2402 | | K+ | | Kaurenoic acid | | Diterpenoid (related to GA) | |  | |
|  | s18 vs sHK | 247.2 | | 247.117 | | 224.1412 | | Na+ | | Methyl jasmonate | | Hormone (JA) | | Yes | |
|  | s18 vs sHK | 267.2 | | 267.102 | | 228.2089 | | K+ | | Myristic acid | | Saturated fatty acids | |  | |
|  |  |  | |  | | 244.2263 | | Na+ | | N1-acetylspermine | | Amino acid | |  | |
|  | s18 vs sHK | 271.2 | | 271.167 | | 248.1412 | | Na+ | | Abscisic acid aldehyde | | Hormone (ABA) | | Yes | |
|  | s18 vs sHK | 686.4 | | 686.391 | | 663.3748 | | Na+ | | 1-Palmitoyl-2-(5-keto-6-octenedioyl)-sn-glycero-3-phosphocholine | | Glycerophospholipids | | Yes | |
| **sHK** | s18 vs sHK | 220.2 | | 220.153 | | 219.1107 | | H+ | | Pantothenate | | Vitamin (B5) | |  | |
|  |  |  | |  | | 219.1120 | | H+ | | Zeatin | | Hormone (cytokinin) | |  | |
|  | s18 vs sHK | 238 | | 238.053 | | 199.0246 | | K+ | | O-phospho-L-homoserine | | Amino acid | |  | |
|  |  |  | |  | | 215.0195 | | Na+ | | O-phospho-4-hydroxy-L-threonine | | Vitamin (B6) | |  | |
|  |  |  | |  | | 215.0807 | | Na+ | | Kinetin | | Hormone (cytokinin) | |  | |
|  | s18 vs sHK | 241.2 | | 241.188 | | 202.2157 | | K+ | | Spermine | | Amino acid | |  | |
|  | s18 vs sHK | 335.2 | | 335.115 | | 334.2144 | | H+ | | Prostaglandin | | Fatty acyls | |  | |
|  |  |  | |  | | 312.3028 | | Na+ | | Eicosanoic acid | | Saturated fatty acids | |  | |
| **sHK** | sHA vs sHK | 355 | | 355.048 | | 354.0577 | | H+ | | 5-amino-6-(5'-phosphoribosylamino)uracil | | Riboflavin | |  | |
| **Strain Associated**  **Table S3. Symbiont-genotype specific metabolites in the high light within the 186b *P. bursaria* host.** These metabolite IDs were highlighted by the pairwise contrasts and were found to have significantly higher abundances in one symbiont-genotype compared to another within the 186b host subset of the *Chlorella* metabolic fraction in the highest light condition (50µE). | **Comparison** | | **mz ID** | | **Detected Mass** | | **Accurate Mass** | | **Adduct** | | **Compound** | | **Pathway** | | **Stress Associated** |
| **s18** | s18 vs sHA | | 171 | | 171.088 | | 132.0059 | | K+ | | Oxalacetic acid | | TCA /central | |  |
|  |  | |  | |  | | 169.9980 | | H+ | | Glycerone phosphate | | Glycolysis / central | |  |
|  |  | |  | |  | | 169.9980 | | H+ | | Glyceraldehyde-3-phosphate | | Glycolysis / central | |  |
|  |  | |  | |  | | 132.0423 | | K+ | | 3-hydroxy-3-methyl-2-oxobutanoate | | Amino acid | |  |
|  |  | |  | |  | | 132.0423 | | K+ | | 2-acetolactate | | Amino acid | |  |
|  |  | |  | |  | | 132.0423 | | K+ | | Glutarate | | Amino acid | |  |
|  |  | |  | |  | | 132.0535 | | K+ | | Asparagine | | Amino acid | |  |
|  |  | |  | |  | | 148.0372 | | Na+ | | Citramalate | | C5-Branched dibasic acid | |  |
|  |  | |  | |  | | 132.0899 | | K+ | | Ornithine | | Amino acid | |  |
|  |  | |  | |  | | 148.0736 | | Na+ | | Mevalonic acid | | Mevalonate pathway | |  |
|  |  | |  | |  | | 148.0736 | | Na+ | | Pantoate | | Pantothenate biosynthesis | |  |
|  | s18 vs sHA | | 237.2 | | 237.181 | | 214.1317 | | Na+ | | Dethiobiotin | | Vitamin (B7) | |  |
|  | s18 vs sHA | | 239.2 | | 239.145 | | 200.1776 | | K+ | | Lauric acid | | Saturated fatty acids | |  |
|  |  | |  | |  | | 216.1725 | | Na+ | | w-hydroxydodecanoic acid | | Hydroxy fatty acids | |  |
|  | s18 vs sHA | | 251.2 | | 251.146 | | 228.2089 | | Na+ | | Myristic acid | | Saturated fatty acids | |  |
|  |  | |  | |  | | 212.2504 | | K+ | | Pentadecane | | Hydrocarbon | |  |
|  | s18 vs sHA | | 537.4 | | 537.356 | | 536.4382 | | H+ | | α/β/γ/δ carotene | | Carotenoid | |  |
|  |  | |  | |  | | 536.4382 | | H+ | | Lycopene (all-trans or tetra cis) | | Carotenoid | |  |
|  | s18 vs sHK + sHA | | 213 | | 213.097 | | 174.0164 | | K+ | | Aconitic acid | | TCA cycle / central | |  |
|  |  | |  | |  | | 190.0114 | | Na+ | | Oxalosuccinate | | TCA cycle / central | |  |
|  |  | |  | |  | | 174.0528 | | K+ | | 3-Carboxy-4-methyl-2-oxopentanoate | | Amino acid | |  |
|  |  | |  | |  | | 174.0528 | | K+ | | Shikimic acid | | Shikimate pathway | |  |
|  |  | |  | |  | | 190.0477 | | Na+ | | 3-dehydroquinate | | Shikimate pathway | |  |
|  |  | |  | |  | | 174.0793 | | K+ | | Indole-3-acetamide | | Amino acid + hormone | |  |
|  |  | |  | |  | | 174.0892 | | K+ | | Suberic acid | | Fatty acid | |  |
|  |  | |  | |  | | 174.1004 | | K+ | | N2-acetyl-L-ornithine | | Amino acid | |  |
|  |  | |  | |  | | 190.1066 | | Na+ | | y-hydroxy-l-arginine | | Arginine-nitric oxide | |  |
|  |  | |  | |  | | 212.0896 | | H+ | | Volemitol | | Carbohydrate | |  |
| *Table S3 continued* | | |  | |  | |  | |  | |  | |  | |  |
| **Strain Associated** | **Comparison** | | **mz ID** | | **Detected mass** | | **Accurate mass** | | **Adduct** | | **Compound** | | **Pathway** | | **Stress Associated** |
|  | s18 vs sHK + sHA | | 257.2 | | 257.123 | | 256.2402 | | H+ | | palmitic acid | | saturated fatty acid | |  |
|  |  | |  | |  | | 256.1172 | | H+ | | 2-(3-Carboxy-3-aminopropyl)-L-histidine | | unusual amino acid | |  |
|  | s18 vs sHK | | 235.2 | | 235.131 | | 212.2504 | | Na+ | | pentadecane | | Hydrocarbon - metabolite | |  |
| **sHK** | s18 vs sHK | | 220.2 | | 220.153 | | 219.1120 | | H+ | | Zeatin | | Hormone | |  |
|  |  | |  | |  | | 219.1107 | | H+ | | Pantothenate | | Vitamin B5 | |  |
| **sHK + sHA** | s18 vs sHK + sHK | | 465 | | 465.096 | | 426.0879 | | K+ | | S-Glutathionyl-L-cysteine | | Cysteine + methionine | | Yes |
| **sHK** | sHA vs sHK | | 329.2 | | 329.1783 | | 328.2402 | | H+ | | Docosahexaenoic acid | | Unsaturated fatty acids | |  |

**Table S4.** **Identified metabolites associated with PCA trajectories for the *P. bursaria* fraction in the evolution experiment.** These were identified from the top 1% of loadings when using the first three principal components. The metabolite ID is that referred to in Figure 5.

| **PC of loading** | **ID** | **Detected**  **mass** | **Accurate**  **mass** | **Adduct** | **Function** | **Pathway** | **Compound** | **Kegg /**  **Metacyc** |
| --- | --- | --- | --- | --- | --- | --- | --- | --- |
| PC1, PC3 | X110.14 | 110 | 109.0197 | H+ | Amino acid | Taurine metab | Hypotaurine | C00519 |
| PC1 | X170.407 | 170 | 131.0582 | K+ | Amino acid + Heme | Heme biosynthesis | 5-Amino-4-oxopentanoate | C00430 |
|  |  |  | 147.0532 | Na+ |  | Amino acid/Central | Glutamate | C00025 |
|  |  |  | 131.0582 | K+ |  | Amino acid | Glutamate 5-semialdehyde | C01165 |
| PC1, PC2, PC3 | X265.213 | 265 | 226.0477 | K+ | Amino acid | Shikimate pathway | Chorismate | C00251 |
|  |  |  | 226.0477 | K+ |  | Shikimate pathway | Prephenate | C00254 |
|  |  |  | 242.0192 | Na+ |  | Shikimate pathway | Deoxy-ketofructose-phosphate | C16848 |
| PC1, PC2, PC3 | X376.2.241 | 376.2 | 353.1699 | Na+ | Plant hormone | Plant hormone (zeatin) | Dihydrozeatin riboside | C16447 |
| PC1, PC2, PC3 | X628.3.437 | 628.3 | 627.2487 | H+ | Plant degradation | Chitin degradation | Chitotriose | CPD-13227 |
| PC1, PC2 | X629.3.437 | 629.3 | 628.2897 | H+ | Plant degradation | Chlorophyll degradation | Chlorophyll catabolite | C18098 |
| PC1, PC2 | X664.3.420 | 664.3 | 625.3033 | K+ | Lipid | Lipid - Arachidonic acid | Leukotriene C4 | C02166 |
| PC1, PC2, PC3 | X685.3.422 | 685.3 | 684.3178 | H+ | Antibiotic | Antibiotic | gamma-L-Glutamyl-butirosin B | C18005 |
| PC1, PC2 | X686.3.421 | 686.3 | 685.3256 | H+ | Antibiotic | Antibiotic | Viomycin | C01540 |
| PC1 | X737.4.422 | 737.4 | 714.3979 | Na+ | Antibiotic | Antibiotic | Avermectin B1b monosaccharide | C11965 |
| PC1, PC3 | X803.4.369 | 803.4 | 780.3622 | Na+ | Antioxidant | Glutathione metabolite | Bis(glutathionyl)spermine | C16563 |
| PC2 | X122.403 | 122 | 121.0197 | H+ | Amino acid | Amino acid | Cysteine | C00736 |
| PC3 | X127.124 | 127 | 88.0160 | K+ | Central | TCA/Glycolysis | Pyruvate | C00022 |
|  |  |  | 104.0110 | Na+ |  | Amino acid | Hydroxypyruvate | C00168 |
| PC3 | X171.66 | 171 | 132.0059 | K+ | Central | Central/TCA/Glycolysis | Oxaloacetate | C00036 |
|  |  |  | 169.9980 | H+ |  | Glycolysis/Carbohydrate | Glycerone phosphate | C00111 |
|  |  |  | 169.9980 | H+ |  | Glycolysis/Carbohydrate | Glyceraldehyde 3-phosphate | C00118 |
|  |  |  | 132.0535 | K+ |  | Amino acid | L-Asparagine | C00152 |
| PC3 | X185.112 | 185 | 146.0215 | K+ | Central | Central/TCA/amino acids | 2-Oxoglutarate | C00026 |
|  |  |  | 146.0579 | K+ |  | Pantothenate + CoA | 2-Dehydropantoate | C00966 |
|  |  |  | 146.0579 | K+ |  | Amino acid | 2-Aceto-2-hydroxybutanoate | C06006 |
| PC3 | X205.124 | 205 | 182.0579 | Na+ | Antioxidant | Amino acid/antioxidant | 4-Hydroxyphenyllactate | C03672 |
|  |  |  | 182.0215 | Na+ |  | Antibiotic | 3;5-Dihydroxyphenylglyoxylate | C12325 |
|  |  |  | 166.0491 | K+ |  | Purine alkaloid | Methylxanthine | C16353 |

| *Table S4 continued* | | | | | | | | |
| --- | --- | --- | --- | --- | --- | --- | --- | --- |
| **PC of loading** | **ID** | **Detected**  **mass** | **Accurate**  **mass** | **Adduct** | **Function** | **Pathway** | **Compound** | **Kegg /**  **Metacyc** |
| PC3 | X220.282 | 220 | 181.0739 | K+ | Amino acid | Amino acid | Tyrosine | C00082 |
|  |  |  | 181.0739 | K+ |  | Amino acid | N-Hydroxy-L-phenylalanine | C19712 |
| PC3 | X289.1.279 | 289.1 | 288.0998 | H+ | Antibiotic | Antibiotic | 6-Deoxydihydrokalafungin | C12435 |
| PC3 | X377.2.241 | 377.2 | 354.2406 | Na+ | Lipid | Lipid - Arachidonic acid | Amoglandin | C00639 |
| PC3 | X393.2.241 | 393.2 | 354.2406 | K+ | Lipid | Lipid - Arachidonic acid | Amoglandin | C00639 |
|  |  |  | 370.2355 | Na+ |  | Lipid - Arachidonic acid | 6-Keto-prostaglandin F1alpha | C05961 |
|  |  |  | 370.2355 | Na+ |  | Lipid - Arachidonic acid | Thromboxane B2 | C05963 |
| PC3 | X398.1.241 | 398.1 | 359.1151 | K+ | Antibiotic | Antibiotic | Penicillin N | C06564 |
|  |  |  | 397.0798 | K+ |  | Antibiotic | 4-Ketoanhydrotetracycline | C06627 |

**Table S5.** **Identified metabolites associated with PCA trajectories for the *Chlorella* fraction in the evolution experiment.** These were identified from the top 1% of loadings when using the first three principal components. The metabolite ID is that referred to in Figure 6.

| **PC of loading** | **ID** | **Detected**  **mass** | **Accurate**  **mass** | **Adduct** | **Function** | **Pathway** | **Compound** | **Kegg /**  **Metacyc** |
| --- | --- | --- | --- | --- | --- | --- | --- | --- |
| PC1, PC2, PC3 | X122.403 | 122 | 121.0197 | H+ | Amino acid | Amino acid | Cysteine | C00097 |
| PC1, PC3 | X393.2.370 | 393.2 | 354.2406 | K+ | Lipid | Arachidonic acid | Amoglandin | C00639 |
|  |  |  | 370.2355 | Na+ |  | Arachidonic acid | 6-Keto-PGF1a | C05961 |
|  |  |  | 370.2355 | Na+ |  | Arachidonic acid | Thromboxane B2 | C05963 |
| PC1, PC2 | X421.2.241 | 421.2 | 382.2508 | K+ | Carotenoid + ABA | Carotenoid + ABA synthesis | C25-Allenic-apo-aldehyde | C14044 |
| PC1, PC3 | X628.3.438 | 628.3 | 627.2487 | H+ | Cell wall metab | Chitin degradation | Chitotriose | CPD-13227 |
| PC1, PC3 | X629.3.437 | 629.3 | 628.2897 | H+ | Chlorophyll degradation | Chlorophyll degradation | Chlorophyll catabolite | C18098 |
| PC1, PC2, PC3 | X664.3.422 | 664.3 | 625.3033 | K+ | Lipid | Arachidonic acid | Leukotriene C4 | C02166 |
| PC2, PC3 | X376.2.241 | 376.2 | 353.1699 | Na+ | Plant hormone | Plant hormone (Zeatin) | Dihydrozeatin riboside | C16447 |
| PC2 | X607.3.420 | 607.3 | 568.305 | K+ | Chlorophyll | Chlorophyll metabolism | Protoporphyrinogen IX | C01079 |
|  |  |  | 584.2635 | Na+ |  | Chlorophyll metabolism | Bilirubin | C00486 |
| PC3 | X217.35 | 217 | 178.0477 | K+ | Secondary metabolite | Ascorbate/Vitamin C | L-Galactono-1;4-lactone | C01115 |
|  |  |  | 194.0579 | Na+ |  | Phenylpropanoid/cell walls | Ferulate | C01494 |
|  |  |  | 216.0399 | H+ |  | Isoprenoid biosynthesis | 2-Methylerythritol 4-phosphate | C11434 |
|  |  |  | 178.063 | K+ |  | Phenylpropanoid/cell wall | Coniferaldehyde | C02666 |
| PC3 | X299.432 | 299 | 260.0297 | K+ | Monosaccharide  phosphate | Starch + sucrose | Glucose 6-phosphate | C00092 |
|  |  |  | 260.0297 | K+ |  | Glycolysis | Glucose 1-phosphate | C00103 |
|  |  |  | 260.0297 | K+ |  | Fructose and mannose | Mannose 6-phosphate | C00275 |
|  |  |  | 276.0246 | Na+ |  | Pentose phosphate | 6-Phospho-D-gluconate | C00345 |
|  |  |  | 260.0297 | K+ |  | Galactose | Galactose 1-phosphate | C00446 |
|  |  |  | 260.0297 | K+ |  | Fructose and mannose | Mannose 1-phosphate | C00636 |

**Table S6.** **Change in symbiont load for each HK1 replicate between the start and end of the evolution experiment**. The metabolic group column denotes whether the replicate’s metabolic profile converged with the profile of the native 186b symbionts or diverged. From these two groups (‘converge’ or ‘diverge’) a group mean difference in symbiont load was calculated.

| **HK1 replicate** | **Difference in symbiont load** | **Metabolic group** | **Group mean**  **difference** |
| --- | --- | --- | --- |
| 1 | 145404.2 | converge |  |
| 2 | 337137.9 | converge | 241271 |
| 3 | 745804.2 | diverge |  |
| 4 | 426775.4 | diverge |  |
| 5 | 490066.7 | diverge | 500951.4 |
| 6 | 341159.3 | diverge | |
